# Sulfatase 2 Inhibition Sensitizes Triple-Negative Breast Cancer Cells to Chemotherapy Through Augmentation of Extracellular ATP

**DOI:** 10.1101/2023.09.15.557965

**Authors:** Jasmine M Manouchehri, Lynn Marcho, Mathew A Cherian

## Abstract

**Background:** Breast cancer is the leading cause of cancer-related death among women worldwide. Patients diagnosed with triple-negative breast cancer (TNBC) have limited therapeutic options that produce durable responses. Hence, a diagnosis of TNBC is associated with a poor prognosis compared to other types of breast cancer. As a result, there is a critical need for novel therapies that can deepen and prolong responses.

We previously found that chemotherapy causes the release of extracellular adenosine triphosphate (eATP). Augmenting eATP release can boost the response of TNBC cells to chemotherapy and cause increased cell death. However, eATP concentrations are limited by several families of extracellular ATPases, which complicates the design of compounds that attenuate eATP degradation.

In this study, we hypothesized that heparan sulfate (HS) would inhibit extracellular ATPases and accentuate chemotherapy-induced cytotoxicity in TNBC by augmenting eATP. HS can be desulfated by sulfatase 1 and 2; sulfatase 2 is consistently highly expressed in a variety of cancers including breast cancer, whereas sulfatase 1 is not. We hypothesized that the sulfatase 2 inhibitor OKN-007 would exacerbate chemotherapy-induced eATP release and TNBC cell death.

**Methods:** TNBC cell lines and nontumorigenic immortal mammary epithelial cells were treated with paclitaxel in the presence of heparan sodium sulfate and/or OKN-007; eATP content and cell viability were evaluated. In addition, protein and cell surface expression of sulfatases 1 and 2 were determined in all examined cell lines via ELISA, Western blot, and flow cytometry analyses.

**Results:** Sulfatase 2 was highly expressed in TNBC cell lines and human breast cancer samples but not in immortal mammary epithelial cells and much less so in normal human breast tissue and ductal carcinoma in situ samples. OKN-007 exacerbated chemotherapy-induced eATP release and chemotherapy-induced TNBC cell death. When combined with chemotherapy, OKN-007 attenuated cells with a cancer-initiating cell phenotype.

**Conclusions:** These results suggest that sulfatase 2 inhibitors in combination with chemotherapy attenuate the viability of TNBC cells more than chemotherapy alone by exacerbating eATP release. These effects, as well as their capacity to attenuate the cancer-initiating cell fraction, may translate into combination therapies for TNBC that induce deeper and more durable responses.

## BACKGROUND

Breast cancer diagnoses impacted 2,261,419 women in 2020 [1]. Among women, this form of cancer has the highest mortality and incidence rates [2]. The mortality rate is largely due to the most aggressive form of breast cancer, triple-negative breast cancer (TNBC). Due to a lack of effective targeted therapies, chemotherapy is still the most efficacious treatment for TNBC, but a major downside is an inability to fully eradicate metastatic disease despite transient responses [3]. Moreover, immunotherapy is only effective in a small subset of patients. Hence, there is a critical need for innovative therapeutic strategies.

The concentration of extracellular adenosine triphosphate (eATP) in tissues is between 0-10 nanomolar (nM) under physiological conditions while the concentration of adenosine triphosphate (ATP) intracellularly ranges from 3-10 millimolar (mM), a more than 10^6^-fold difference [4]. However, this minute eATP quantity fulfills a critical role as a signaling molecule through cell surface purinergic receptors. Moreover, there is a marked increase in the concentration of eATP in the tumor microenvironment (TME), which can reach the micromolar range [5–7]. Our published study showed that eATP is toxic to TNBC cells in the high micromolar range but not in nontumorigenic immortal mammary epithelial MCF-10A cells [8]. Thus, cancer cells “live” closer to the threshold for cytotoxicity.

Furthermore, our published data showed that chemotherapy treatment results in augmentation of eATP concentration [8]. However, eATP concentration is limited by several families of ecto-nucleotidases, including ecto-nucleoside triphosphate diphosphohydrolases (ENTPD), 5’-nucleotidases (5’-NTs), ecto-nucleotide pyrophosphatases/phosphodiesterases (E-NPPase), and tissue nonspecific alkaline phosphatases (TNAP), with E-NTPD considered to be the central enzyme responsible for ATP degradation and extracellular 5’-NT responsible for the catalytic conversion of AMP to adenosine and inorganic phosphate [9]. We previously showed that inhibitors of each of these families of ecto-ATPases can augment chemotherapy-induced TNBC cell death and eATP release through P2RX4 and P2RX7 ion-coupled purinergic receptors. This data suggests that all ecto-ATPases may need to be inhibited to maximize eATP release and TNBC cell death. The existence of several families of structurally diverse ecto-ATPases complicates the design of small-molecule inhibitors that can maximally suppress eATP degradation.

We noted that sulfated polysaccharides have been reported to inhibit multiple families of extracellular ATPases with nanomolar potency; the degree of sulfation, which imparts a negative charge to the molecule, is critical for this inhibition [10]. We also noted that endogenous extracellular polysulfated polysaccharide heparan sulfate (HS) inhibits eATP degradation [11]. Moreover, other publications show that the ecto-ATPase ecto-nucleotide pyrophosphatases/phosphodiesterase 1 (E-NPP1) binds to extracellular glycosaminoglycans, and its ATPase activity is competitively inhibited by these glycosaminoglycans, including heparin and HS [12].

HS proteoglycans are ubiquitously expressed in the extracellular matrix and cell surface of animal cells. They can be broadly divided into three groups: transmembrane syndecans; glycosylphosphatidylinositol-linked glypicans; and extracellular matrix-associated perlecan, agrin, and collagen XVIII [13]. The polysulfated polysaccharide HS is synthesized in the Golgi system and is composed of disaccharide units that are negatively charged and unbranched with sulfation of the 3-O, 6-O, or N sites of glucosamine along with the 6-O site of glucuronic/iduronic acid [14–17]. HS plays a tumor suppressor role: loss of exostosin 1 or exostosin 2, proteins involved in the polymerization of HS, leads to hereditary multiple exostoses, a hereditary cancer syndrome associated with an elevated risk of chondrosarcomas and osteosarcomas [18–21]. Sulfatase 1 and sulfatase 2 are extracellular HS 6-O-endosulfatases that hydrolyze the 6-O-sulfate groups on glucosamine residues in HS [14, 22–25]. Sulfatase 2 is highly expressed in a variety of cancers including breast cancer, while sulfatase 1 is not [15, 22, 25–27]. Sulfatase 2 has been shown to enhance tumor initiation and progression in a variety of cancers including breast **(Supplemental Figure 1)** [24, 26, 28]. When sulfatase 2 was overexpressed in the breast cancer cell line MDA-MB 231, there was an increase in breast cancer cell growth [24]. In contrast to sulfatase 2, sulfatase 1 is a tumor suppressor, possibly due to its negative regulation of fibroblast growth factor signaling [22, 29]. Notably, the expression of sulfotransferase 3-OST3A, an enzyme that adds sulfate groups to HS, is epigenetically downregulated in estrogen receptor-positive (ER+) and TNBC cell lines as well as chondrosarcoma cell lines [30, 31]. In addition, expression of sulfotransferase 3-OST3A is epigenetically silenced in human breast, colon, lung, and pancreatic cancers [32, 33].

OKN-007 is a potent sulfatase inhibitor that has been demonstrated to decrease sulfatase 2 activity [34]. Moreover, OKN-007 is already in phase Ib clinical trials for glioblastoma multiforme (NCT03587038). Hence, we hypothesized that sulfatase 2 inhibitors would accentuate the fully sulfated form of HS in the TME, an endogenous inhibitor of ecto-ATPases, thus enhancing TNBC cell death by augmenting eATP concentrations in the microenvironment of chemotherapy-treated cells **(Figure 1)**.

**Figure 1.**
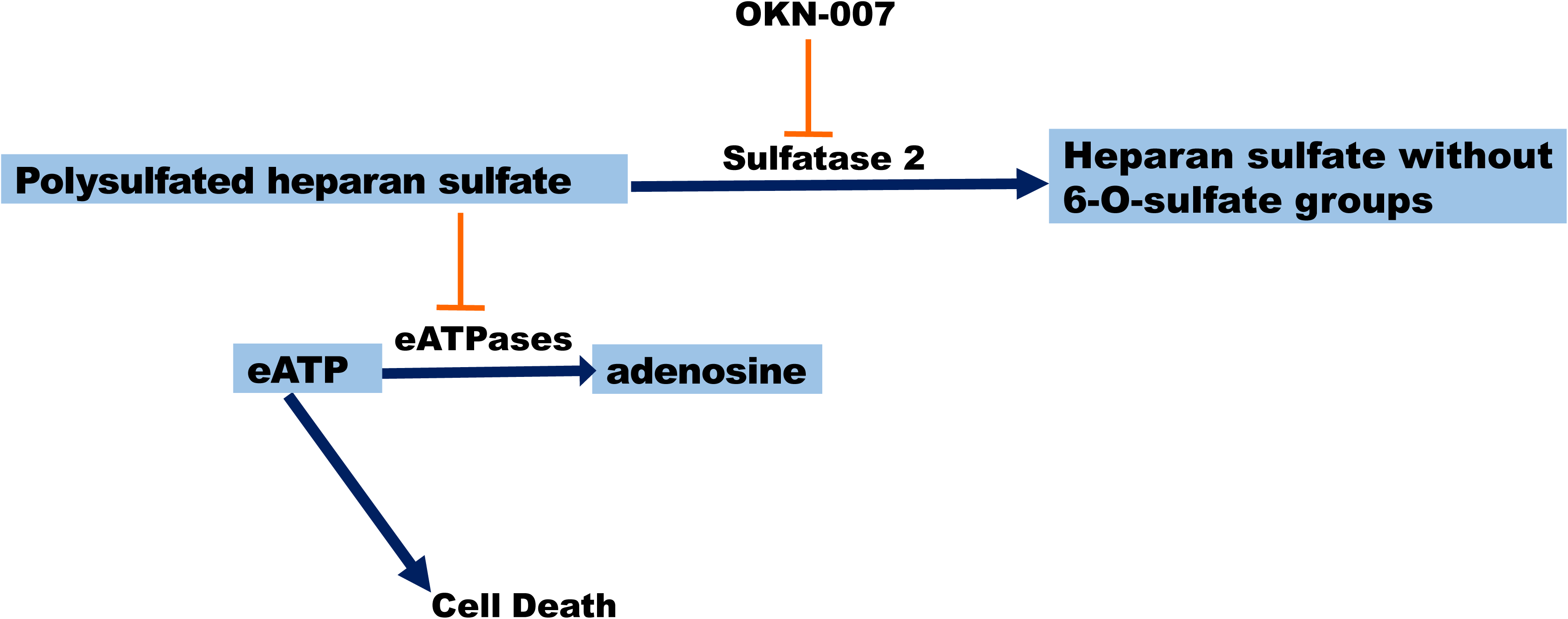
Schematic displaying our proposed model for heparan sulfate’s impact on eATP. Our proposed model suggests that accumulation of polysulfated polysaccharide HS, due to the presence of the sulfatase inhibitor OKN-007, facilitates ATP accumulation in the extracellular environment of paclitaxel-treated TNBC cells by preventing the breakdown of eATP by eATPases, which can lead to exacerbation of chemotherapy-induced cell death.

## METHODS

### Cell culture, drugs, and chemicals

Breast cancer cell lines MDA-MB 231 (ATCC HTB-26, RRID: CVCL_0062), MDA-MB 468 (ATCC HTB-132, RRID: CVCL_0419), Hs 578t (ATCC HTB-126, RRID: CVCL_0332), HEK-293T ATCC Cat# CRL-3216, RRID: CVCL_0063), and nontumorigenic immortal mammary epithelial MCF-10A cells (ATCC Cat# CRL-10317, RRID: CVCL_0598) were authenticated and maintained as described previously [8].

The following drugs and chemicals were used: ATP (Sigma), dimethyl sulfoxide/DMSO (Sigma), paclitaxel (Calbiochem), OKN-007 (formally known as NXY-059) (Selleck Chemical), and heparan sodium sulfate (Sigma). HS was dissolved in nuclease-free water (Invitrogen); paclitaxel and OKN-007 were dissolved in dimethyl sulfoxide (DMSO) (Sigma). **Table 1** shows the drugs’ concentrations and functions; we optimized the drug concentrations that were applied for the different assays by using previously described drug concentrations as starting points [34–38]. Cells were treated at the designated concentrations.

**Table 1.** Drug concentrations and functions.

| Drug | Concentration(s) | Function | Concentration reference |
| --- | --- | --- | --- |
| paclitaxel | 50 and 100 μM | Chemotherapeutic agent | [1] |
| OKN-007 | 20 μM | Sulfatase inhibitor | [2] |
| A437809 | 20 μM | P2RX7 inhibitor | [3] |
| 5-BDBD | 20 μM | P2RX4 inhibitor | [4] |
| heparan sodium sulfate | 50 μM | developmental processes, angiogenesis, and tumor metastasis | [5] |

### Measurement of ecto ATPase inhibition by heparan sulfate

ATP was combined with recombinant proteins (enzymes): ENPP1, tissue non-specific alkaline phosphatase (TNAP), and ecto-nucleoside triphosphate diphosphohydrolases 1 (CD39/ENTPD1) (R&D Systems) in the presence or absence of heparan sodium sulfate. The amount of eATP was analyzed according to the protocol described by the manufacturer of ATPlite 1-step Luminescence Assay System (PerkinElmer, Cat# 6016736) either at 0 hours (eATP concentrations were determined immediately) or after 24 hours of incubation at 37°C. Luminescence readings were obtained from a Bioteck Synergy HT plate reader. As a control, enzymes were boiled, and the subsequent experiments were carried out in the same fashion. The student’s t-test was applied to ascertain significance. **represents p <0.01 for the comparison of enzyme-treated ATP to enzyme-treated ATP and heparan sodium sulfate.

### Western blot analysis of expression of sulfatases

An equal number of individual cell lines (TNBC MDA-MB 231, Hs 578t, MDA-MB 468 cells and nontumorigenic immortal mammary epithelial MCF-10A cells) were seeded and cultured for 48 hours to 70%-80% confluency. Cell supernatants were collected. Total cell lysates were prepared; protein quantification was performed; and proteins were denatured, separated, and transferred as previously described [8]. For immunoblot of sulfatase 1 and sulfatase 2 on cell lysates, 100 µg of protein was loaded onto gels. For immunoblot of sulfatase 1 and sulfatase 2 on cell supernatants, either 10 µL of cell supernatants for the analysis of unnormalized expression levels or a volume that was normalized to the total mass of protein measured in the corresponding lysate was loaded onto the gels. Unadjusted (with 10 µl supernatant) blots were stained with Ponceau S (Thermo Fisher Scientific) as a loading control. The membranes were blocked with nonfat milk at room temperature for an hour and incubated overnight at 4°C with a primary antibody: sulfatase 1 (1:200 dilution; Novus Biologicals, Cat# NBP231584, RRID: AB_2916043) and sulfatase 2 (1:500 dilution; Abcam, Cat# ab232835, RRID: AB_2916044) diluted in 5% nonfat milk. The membranes were washed and developed as described previously [8].

Glyceraldehyde-3-phosphate dehydrogenase (GAPDH) (Cell Signaling Technology, Cat #3683, RRID: AB_1642205) was used as a loading control. Densitometry was performed on Licor Image Studio (RRID: SCR_015795). The student’s t-test was used for the applicable assays to ascertain significance. * represents p<0.05 and ** represents p<0.01 relative to protein expression in MCF-10A cells.

### ELISAs

TNBC cell lines, MDA-MB 231, Hs 578t, MDA-MB 468 cells, and nontumorigenic immortal mammary epithelial MCF-10A cells were grown for 48 hours to 70-80% confluency, and supernatants were collected. The concentrations of sulfatase 1 and 2 (Lifespan Biosciences, Cat# LS-F66757-1 and LS-F35926-1) were measured in the supernatants of the examined cell lines at basal level via enzyme-linked immunoassay (ELISA) analysis according to the manufacturer’s directions.

### Flow cytometry analysis of sulfatases

Cell surface expressions for sulfatase 1 and sulfatase 2 were measured in TNBC cell lines, MDA-MB 231, Hs 578t, MDA-MB 468 cells, nontumorigenic immortal mammary epithelial MCF-10A cells, and HEK 293T cells. HEK 293T cells were transfected with either sulfatase 1 or sulfatase 2 expression plasmids or empty vector control derived from pcDNA3.1 (RRID: Addgene_79663) using Lipofectamine 3000 (Thermo Fisher Scientific). Cells were detached with accutase (Thermo Fisher Scientific). One million cells were washed in phosphate buffered saline (PBS) with 0.05% BSA, stained with sulfatase 1 (Novus Biologicals, Cat# NBP231584, RRID: AB_2916043) or sulfatase 2 (Abcam, Cat# ab232835, RRID: AB_2916044) plus goat anti-rabbit IgG (H+L) secondary antibody-fluorescein isothiocyanate (FITC) (Novus Biologicals, Cat# NB 7168, RRID: AB_524413) or stained with rabbit IgG Isotype Control-FITC (Invitrogen, Cat# PA5-23092, RRID: AB_2540619) in flow cytometry staining buffer (2% FBS, 0.02% sodium azide and PBS). Analysis was performed on BD fluorescence-activated cell sorting (FACS) Fortessa using the FITC channel (530/30 nm) and Flowjo software (RRID: SCR_008520). The student’s t-test was applied to ascertain significance. * represents p<0.05 and ** represents p<0.01 relative to mean fluorescence intensity (MFI) in MCF-10A cells; + represents p<0.05 and ++ represents p<0.01 relative to MFI in HEK293-empty vector transfected. O/E represents overexpressed.

### Sulfatase activity assay

The sulfatase inhibitor OKN-007 (20 µM) or vehicle control was added to recombinant sulfatase enzyme (0, 2, 4 µL) with sulfatase substrate (p-nitrocathecol sulfate) diluted in the sulfatase assay buffer, and sulfatase activity was determined by comparison with the p-nitrocathecol standard dilution curve. The p-nitrocathecol standard was prepared according to the manufacturer’s protocol for the sulfatase activity assay kit (Sigma, Cat# MAK276-1KT). Absorbance readings at 515 nm were obtained from a Bioteck Synergy HT plate reader. The student’s t-test was applied to ascertain significance. * represents p<0.05 and ** represents p<0.01, comparing the sulfatase activity between the recombinant sulfatase enzyme and vehicle control addition and the combination of the sulfatase enzyme and OKN-007.

### Immunohistochemistry of sulfatase 2

AMSBIO BR1202B breast cancer tissue array (120-core array) was sectioned at 5 µm and air-dried overnight on Fisher Superfrost Plus slides. Also, normal breast tissue or ductal carcinoma in situ (DCIS) was prepared on PERMAFLEX plus slides. All of the stainings were performed at Histowiz, Inc. (Brooklyn, NY) using the Leica Bond RX automated stainer (Leica Microsystems) and using a standard operating procedure with a fully automated workflow. Normal breast and DCIS samples were processed, embedded in paraffin, and sectioned at 4 μm. The slides were dewaxed using xylene and alcohol-based dewaxing solutions. Immunohistochemistry: Epitope retrieval was performed by heat-induced epitope retrieval (HIER) of the formalin-fixed, paraffin-embedded tissue using citrate-based pH 6 solution (Leica Microsystems, Cat#AR9961) for 10 minutes at 95°C. The tissues were first incubated with a peroxide block buffer (Leica Microsystems, Cat# RE7101-CE), followed by incubation with the rabbit anti-sulfatase 2 antibody (Abcam, Cat# ab232835, RRID: AB_2916044) at 1:50 dilution for 30 minutes, followed by DAB rabbit secondary reagents: polymer, DAB refine, and hematoxylin (Bond Polymer Refine Detection Kit, Leica Microsystems, Cat# DS9800) according to the manufacturer’s protocol. The slides were dried, coverslipped (TissueTek-Prisma Coverslipper), and visualized using a Leica Aperio AT2 slide scanner (Leica Microsystems) at 40×.

Quantitation of protein expression: Automated analysis of protein expression was done at Histowiz, Inc. For the analysis of the tissue microarray (TMA), the Halo TMA module was used to identify and extract the individual TMA cores by means of constructing a grid over the TMA. All subsequent analysis steps were the same for the 3 slides with DCIS tissue and 2 slides with normal breast tissue and the TMA slide. In the first part of the analysis, the tumor area was identified by training a random forest classifier algorithm to separate viable tumor tissue from any surrounding stroma and necrosis areas. Once the tumor area was identified, the analysis then proceeded to identify positive and negative cells based on sulfatase 2 staining within the defined tumor area on each slide and each core from the TMA slide. Positive and negative cells were identified using the Halo Multiplex IHC algorithm v3.4.1 by first defining the settings for the hematoxylin counterstain, followed by setting thresholds to detect the sulfatase 2 stain positivity of weak, moderate, and strong (Halo threshold settings 0.11, 0.35, 0.45). An H-score was then generated following the convention below: weak positive (1+), moderate positive (2+), and strong positive (3+). For the immunohistochemistry statistical analysis, group differences were calculated using Kruskal-Wallis one-way analysis of variance (ANOVA). When shown to be statistically significant, a post hoc Dunn’s test was done to determine p values. P values were adjusted to account for multiple comparisons, and an alpha level of 0.05 was used for all the tests. The software GraphPad Prism version 10.0.1 (RRID: SCR_002798) was used for these tests.

### Cell viability and eATP assays

TNBC MDA-MB 231, Hs 578t, MDA-MB 468 cells, and nontumorigenic immortal mammary epithelial MCF-10A cells were plated as previously described and treated with paclitaxel (vehicle), heparan sodium sulfate (50 µM), OKN-007 (20 µM), A438709 (20 µM), 5-BDBD (20 µM), or different combinations of these drugs. Cells were treated with OKN-007 and heparan sodium sulfate for 48 hours and with paclitaxel, A438709, or 5-BDBD for the final 6 hours of the 48-hour time course (we treated cells with paclitaxel for 6 hours to replicate exposure times in patients); cell viability was assessed by applying the PrestoBlue™ HS cell viability reagent (Invitrogen, Cat# P50201) following the manufacturer’s instructions [8]. ATP was assessed in supernatants as described above. Fluorescence readings (excitation and emission ranges: 540–570 nm and 580–610 nm) were assessed using a Bioteck Synergy HT plate reader. One-way ANOVA with Tukey’s honestly significant difference (HSD) was calculated to ascertain significance. * represents p<0.05 and ** represents p<0.01 when comparing vehicle addition to OKN-007, to HS, and the combination of HS, OKN-007, and/or paclitaxel.

### Flow cytometry for cancer-initiating cells

TNBC MDA-MB 231, Hs 578t, and MDA-MB 468 cells were collected and stained following the protocol for the Aldeflour Kit (STEMCELL, Cat# 01700). Cells were washed and stained with CD24-PE (eBioscience, Cat# 12-0247-42, RRID: AB_1548678), CD44-APC (eBioscience, Cat# 17-0441-82, RRID: AB_469390), and LIVE/DEAD™ Fixable Near-IR Dead Cell Stain Kit (Invitrogen, Cat# L10119) for 30 minutes at 4°C. Cells were washed and resuspended in the Aldefluor Buffer provided in the Aldefluor kit. Flow cytometry analysis was performed on BD FACS Fortessa using the FITC (ALDH), PE (CD24), APC (CD44), and AF750 (Live/Dead) channels and applying the Flowjo software (RRID: SCR_008520). One-way ANOVA with Tukey’s HSD was calculated to ascertain significance. For statistical analysis, * represents p<0.05 and ** represents p<0.01 when comparing paclitaxel to the combination of paclitaxel and OKN-007.

### Tumorsphere formation efficiency assay

TNBC MDA-MB 231, Hs 578t, and MDA-MB 468 cell lines were grown and treated with paclitaxel, OKN-007, and/or heparan sodium sulfate as described above in the “Cell viability and eATP assay” section. Cells were trypsinized, washed, resuspended in 3D Tumorsphere Medium XF (Sigma), and plated at 10 viable cells per well after (45 µM) filtration on low-attachment, round-bottom 96-well plates. Cells were grown for 7 days, and tumorspheres were counted for each different condition using the Etaluma™ Lumascope 620. One-way ANOVA with Tukey’s HSD was calculated to ascertain significance. For statistical analysis, ** represents p<0.01 when comparing paclitaxel to the combination of paclitaxel and OKN-007.

## RESULTS

### Effect of heparan sulfate on ecto-ATPase activity

We first sought to determine if polysulfated HS prevents eATP degradation by different families of ecto-ATPases: tissue non-specific alkaline phosphatase (TNAP), ecto-nucleoside triphosphate diphosphohydrolases 1 (ENTPD1), and ecto-nucleotide pyrophosphatase/phosphodiesterase (ENPP1). We incubated ATP (500 µM) with vehicle or 100 units of one member of each of the families of ecto-ATPase enzymes in the presence or absence of heparan sodium sulfate (50 µM) for 48 hours or 0 hours (ATP concentrations determined immediately) at 37°C **(Figures 2A and B)**. In the absence of heparan sodium sulfate, all three enzymes significantly decreased the concentration of ATP after 48 hours, but in the presence of heparan sodium sulfate, eATP concentrations were not depressed. Additionally, as a control, we used boiled enzymes added to ATP in the presence or absence of heparan sodium sulfate, and the eATP concentrations did not change, demonstrating that the reduction in ATP observed with the addition of enzymes is specifically due to their catalytic activity **(Figures 2C and D)**. We also did not observe any change in ATP concentration when its level was measured at 0 hours (immediately) after the addition of enzyme. This result indicates that HS does not positively or negatively interfere with the ATP assay. These results demonstrate that HS inhibits the ATPase activity of each of the three major families of ecto-ATPases.

**Figure 2.**
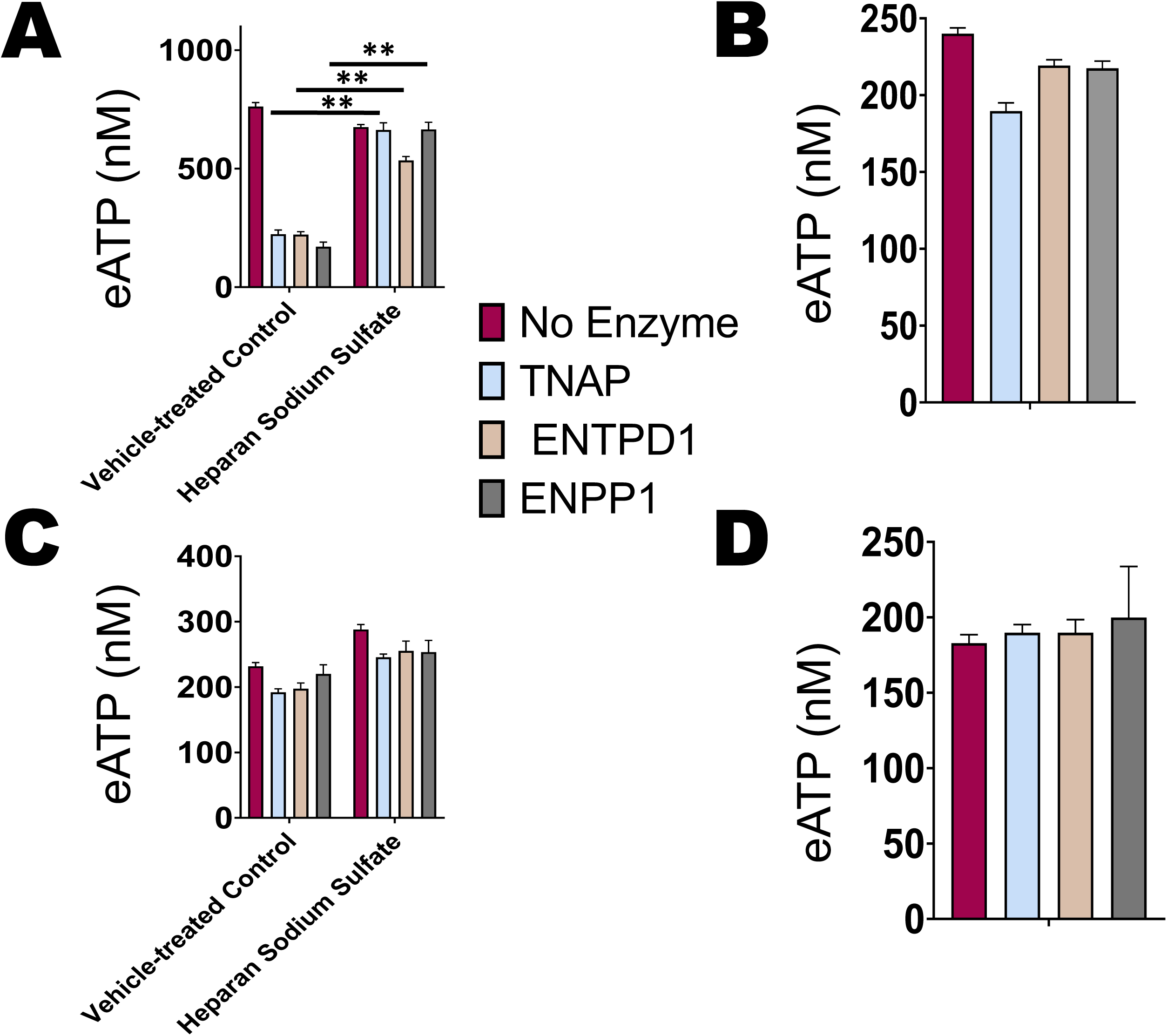
Heparan sulfate’s influence on eATPases’ activity. ATP (500 µM) was incubated with vehicle or 100 units of one member of each of the families of extracellular ATPases, tissue non-specific alkaline phosphatase (TNAP), ENTPD1, and ENPP1, in the presence or absence of heparan sodium sulfate (50 µM) for **(A)** 48 hours at 37°C or **(B)** 0 hours. For **(C)** and **(D)**, enzymes were boiled before incubation, and eATP was measured after **(C)** 48 hours at 37°C or **(D)** 0 hours. ATP concentrations were measured using a luciferase-based assay. HS blocked eATP degradation by all three families of enzymes. The standard deviation was calculated from three independent experiments performed in triplicate. The student’s t-test was performed to determine significance, with ** indicating *p* value <0.01 for the comparison of enzyme-treated ATP vs. enzyme and heparan sodium sulfate.

### Analysis of expression of sulfatases in breast cancer

#### Expression of sulfatases in breast cancer cell lines and mammary epithelial cells by Western blot and ELISA

To compare the level of expression of sulfatase 2 and sulfatase 1 in TNBC cell lines to that in control immortal mammary epithelial cells, we performed Western blot analysis on TNBC MDA-MB 231, Hs 578t, and MDA-MB 468 cell lines and nontumorigenic immortal epithelial mammary MCF-10A cells, probing for sulfatase 2 and sulfatase 1 with GAPDH as the internal loading control **(Figures 3 and 4; Supplemental Figures 2-7)**. TNBC cell lines expressed markedly higher levels of sulfatase 2 protein intracellularly as well as extracellularly in supernatants when compared to MCF-10A cells, as assessed by semi-quantitative densitometry. We separately probed for sulfatase 1 protein in the same cell lines and determined that the TNBC cell lines express more sulfatase 1 extracellularly in comparison to the immortal MCF-10A cells, but Hs578t was the only TNBC cell line to express more intracellular sulfatase 1 protein in comparison to immortal MCF-10A cells. We also checked the baseline expression levels of sulfatases in the supernatants of TNBC cell lines and MCF-10A cells via ELISAs **(Supplemental Figure 8)**. We observed that TNBC cell lines expressed significantly more sulfatase 2 in their supernatants in comparison to MCF-10A cells aligning with the Western blot analysis. TNBC cell lines expressed less sulfatase 1 in their supernatants in comparison to control MCF-10A cells.

**Figure 3.**
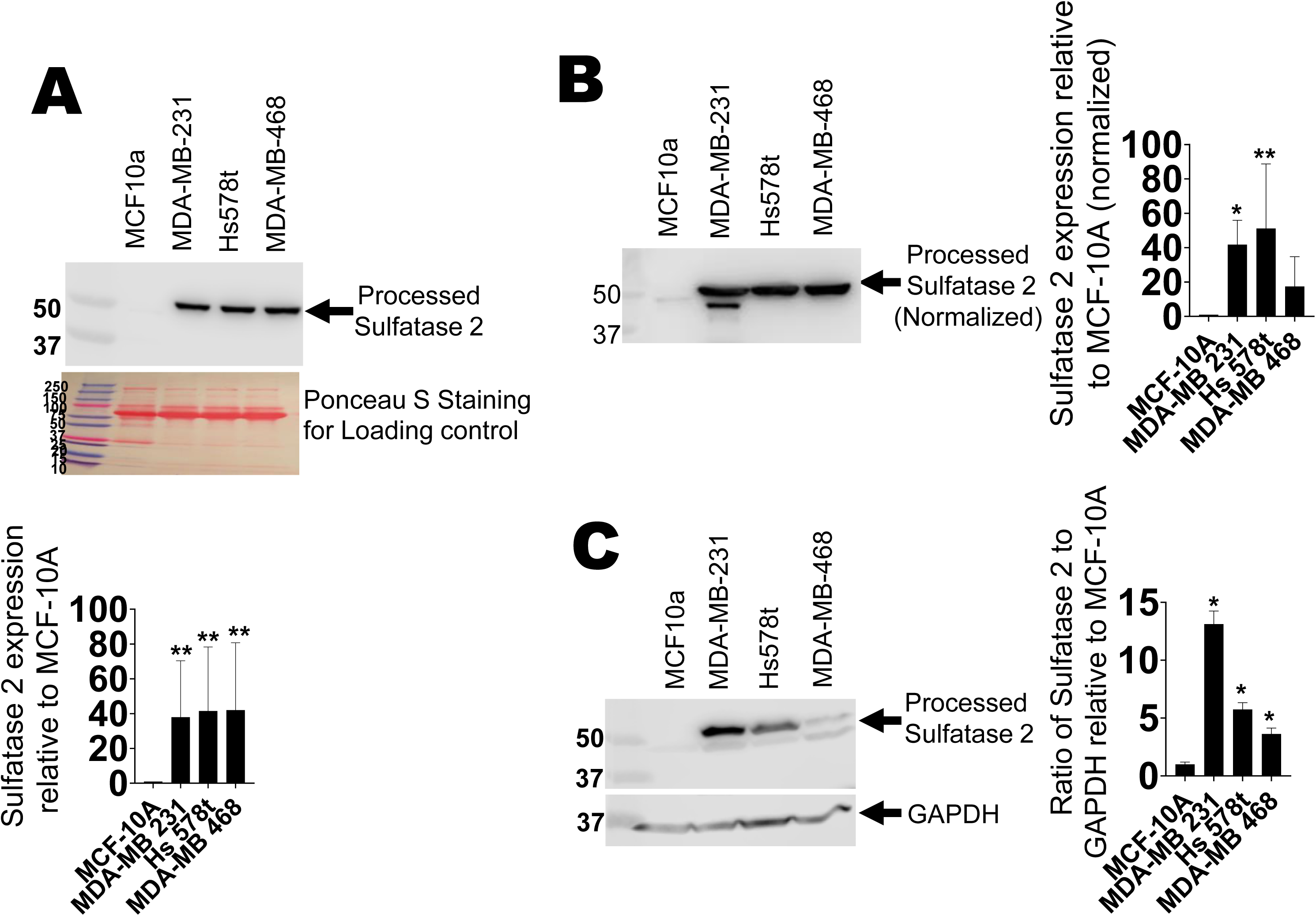
Immunoblot analysis of sulfatase 2 expression in immortal mammary epithelial cells and TNBC cell lines. For the Western blot analysis of sulfatase 2, **(A)** 5 μl of cell supernatants of nontumorigenic immortal mammary epithelial MCF-10A cells and TNBC cell lines—MDA-MB 231, Hs 578t, and MDA-MB 468 cells—were subjected to polyacrylamide gel electrophoresis (PAGE) and probed with a human sulfatase 2-specific antibody. **(B)** Adjusted volume of cell supernatants inversely proportionate to the protein concentration in the corresponding cell lysate were loaded for PAGE and probed with a sulfatase 2-specific antibody to control for cell number and viability. **(C)** Equal amounts of cell lysate from each cell line were probed with a sulfatase 2-specific antibody. All experiments were repeated once with biological replicates. The densitometric analyses of the bands were performed using Image Studio software (LI-COR Inc.). The student’s t-test was performed to determine significance with * representing *p*<0.05 and ** representing *p*<0.01 when comparing expression in MCF-10A cells to expression in TNBC cell lines. Cropped blots are shown in the figure and full-length blots are presented in **Supplementary** Figures 3-5.

**Figure 4.**
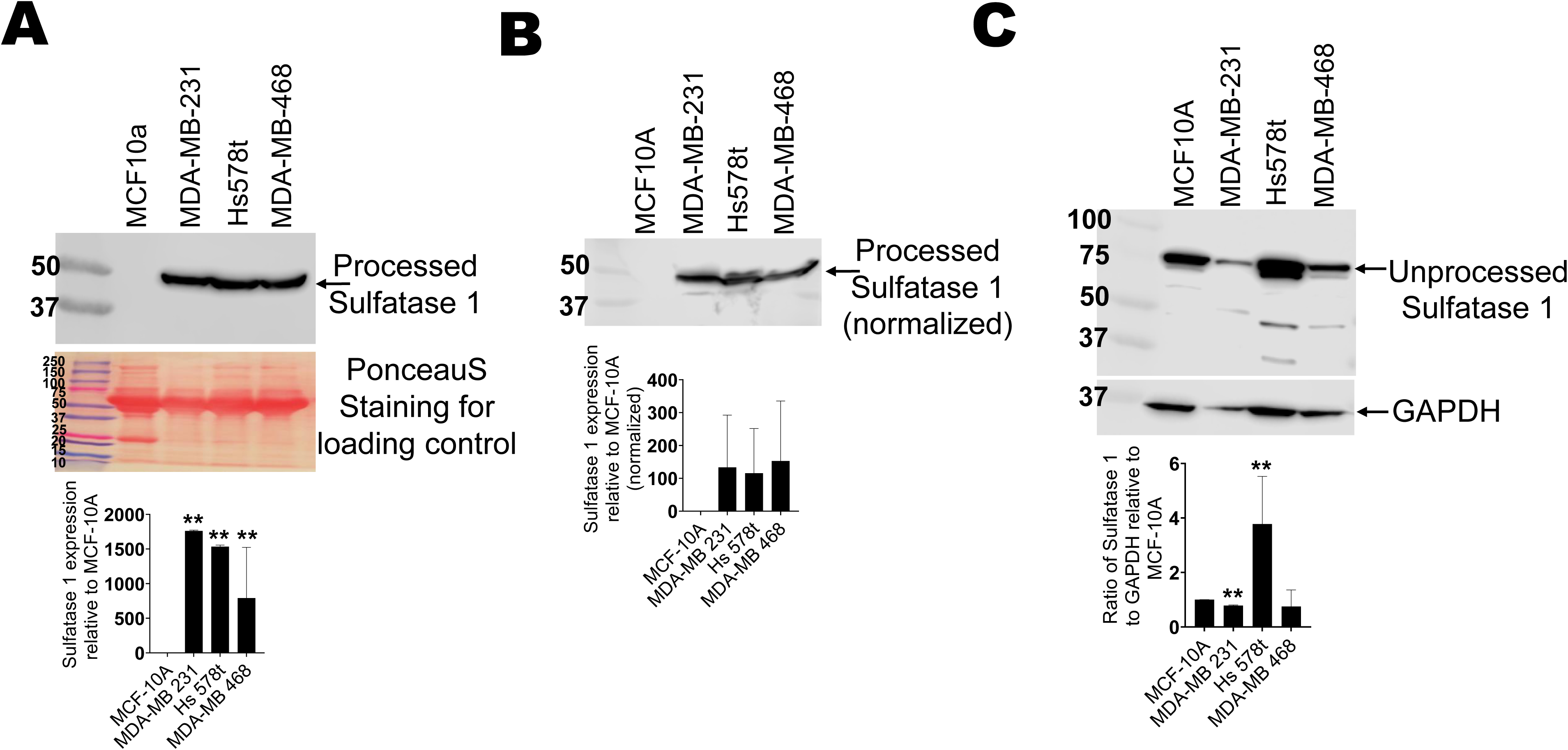
Immunoblot analysis of sulfatase 1 expression in immortal mammary epithelial cells and TNBC cell lines. For the Western blot analysis of sulfatase 1, **(A)** 5 μl of cell supernatants of nontumorigenic immortal mammary epithelial MCF-10A cells and TNBC cell lines—MDA-MB 231, Hs 578t, and MDA-MB 468 cells—were subjected to PAGE and probed with a human sulfatase 1-specific antibody. **(B)** Adjusted volume of cell supernatants inversely proportionate to the protein concentration in the corresponding cell lysate were loaded for PAGE and probed with a sulfatase 1-specific antibody. **(C)** Equal amounts of cell lysate from each cell line were probed with a sulfatase 1-specific antibody. All experiments were repeated once with biological replicates. The densitometric analyses of the bands were performed using Image Studio software (LI-COR Inc.). The student’s t-test was performed to determine significance with * representing *p*<0.05 and ** representing *p*<0.01 when comparing expression in MCF-10A cells to expression in TNBC cell lines. Cropped blots are shown in the figure and full-length blots are presented in **Supplementary** Figures 6-8.

#### Flow cytometry analysis of cell surface expression of sulfatases in breast cancer cell lines and immortal mammary epithelial cells

We examined the cell surface expressions of sulfatase 1 and sulfatase 2 in TNBC cell lines and immortal mammary epithelial cells. We performed flow cytometry analysis on TNBCs, MCF-10A, and HEK 293T cells transfected with sulfatase 2 as a positive control probing for cell surface expression of sulfatase 2 **(Figure 5A; Supplemental Figure 9)**. MDA-MB 231, Hs 578t, and MDA-MB 468 did not express significantly more or less cell surface sulfatase 2 in comparison to control MCF-10A cells. We also probed all the cell lines for sulfatase 1 and observed that immortal MCF-10A expressed the most cell surface sulfatase 1 in comparison to the expressions in TNBC cell lines, which aligns with the results from ELISAs.

**Figure 5.**
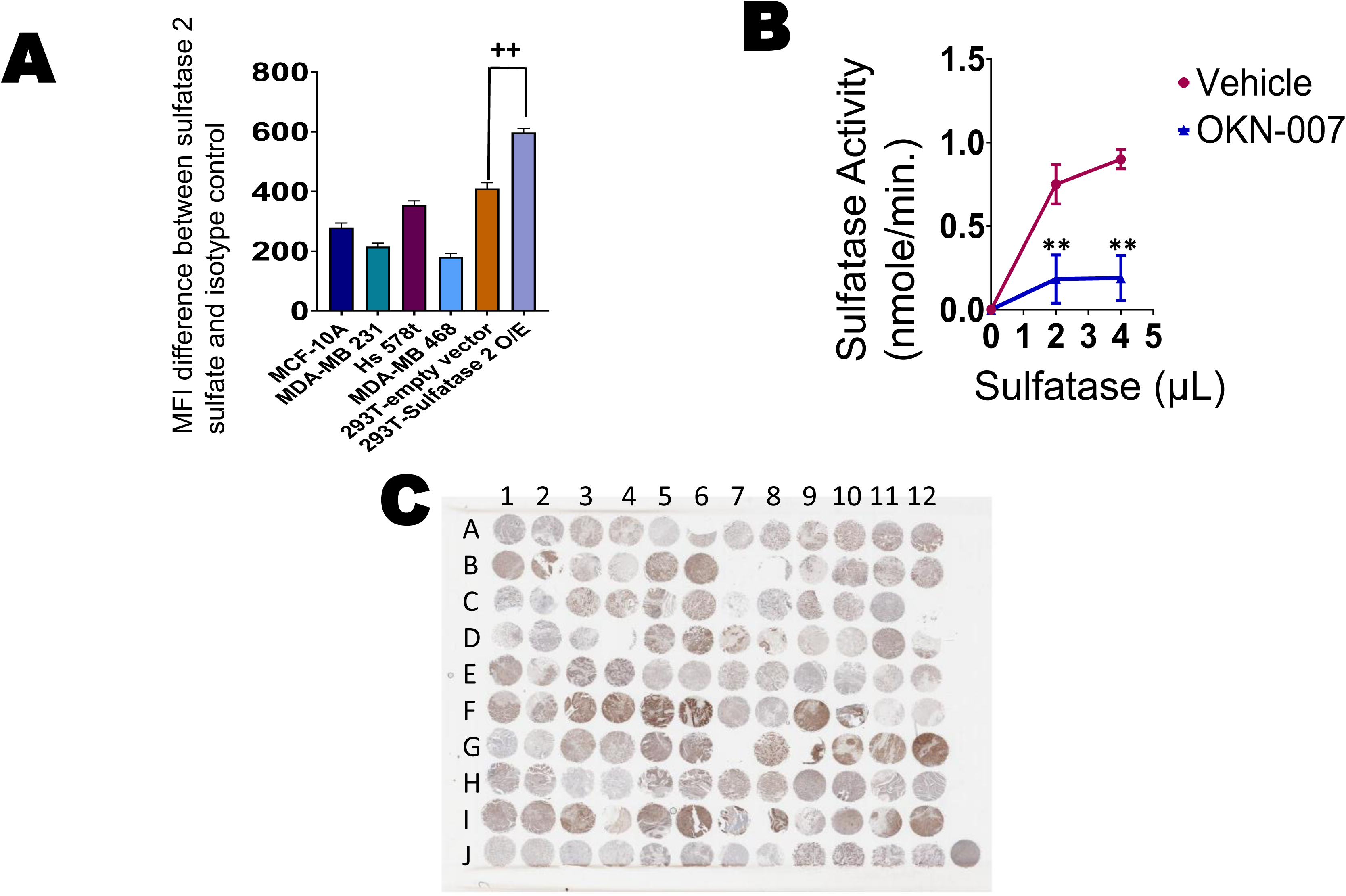
Cell surface expression of sulfatase 2, sulfatase activity, and immunohistochemistry. (**A**) Baseline sulfatase 2 cell surface expression levels were examined via flow cytometry analysis in nontumorigenic immortal mammary epithelial MCF-10A cells and TNBC MDA-MB 231, Hs 578t, and MDA-MB 468 cells by flow cytometry. The standard deviation (SD) was calculated from three independent experiments performed in triplicate. The student’s t-test was performed to determine significance with * representing *p*<0.05 and ** representing *p*<0.01, comparing expression levels in MCF-10A to those in the TNBC cell lines. **(B)** A sulfatase activity assay was carried out. OKN-007 (20 µM) was added to the vehicle (sulfatase; 0, 2, 4 µL) and sulfatase substrate; sulfatase activity (desulfation of p-nitrocathecol sulfate to p-nitrocathecol) was determined using a standard dilution of p-nitrocathecol. There was a decrease in sulfatase activity when sulfatase and sulfatase substrate were exposed to OKN-007, confirming the antagonistic effect of this inhibitor on sulfatase. The SD was calculated from three independent experiments performed in triplicate. The student’s t-test was performed to determine significance with * representing *p*<0.05 and ** representing *p*<0.01, comparing the sulfatase activity between the vehicle (sulfatase) to OKN-007 and vehicle. **(C)** The AMSBIO BR1202B breast cancer tissue array (120 cores with 82 TNBC cores; key can be found in **Supplemental Figure 10C**) was stained with sulfatase 2.

### Effect of OKN-007 on sulfatase activity

We wanted to confirm that the sulfatase 2 inhibitor OKN-007 inhibits sulfatase activity at the concentrations planned for our experiments. We utilized a sulfatase activity kit that can measure the desulfation of p-nitrocathecol sulfate by sulfatase to p-nitrocatechol. We added the sulfatase inhibitor OKN-007 (20 µM) to recombinant sulfatase (0, 2, 4 µL) and sulfatase substrate diluted in the sulfatase assay buffer, and sulfatase activity was determined through a calculated standard dilution of p-nitrocathecol **(Figure 5B)**. We found that there was significantly decreased sulfatase activity when sulfatase and sulfatase substrate were exposed to OKN-007. These results demonstrate that OKN-007 neutralizes sulfatase activity at the concentration utilized (20 µM).

### Measurement of sulfatase 2 expression in human breast cancer samples by immunohistochemistry

An AMSBIO BR1202B breast cancer tissue array (120 cores with 82 TNBC cores) was stained with sulfatase 2 **(Figure 5C; Supplemental Table 1; Supplemental Figure 10)**. We focused on sulfatase 2 because sulfatase 1 has been reported to be a tumor suppressor [24, 39]. Statistical analysis was performed on the breast cancer tissue array stained with sulfatase 2 antibody. The expression of sulfatase 2 was compared between TNBCs, ER+/progesterone receptor-positive (PR+), human epidermal growth factor receptor 2 positive (HER2+) breast cancer, normal breast tissue, and DCIS. We carried out the Kruskal-Wallis test and found no significant difference when: comparing the average percentages of cells expressing any level of sulfatase 2 between the different groups **(Figure 6A)**; the percentages of cells that were weakly positive for sulfatase 2 **(Figure 6B)**; and the average percentages of cells that did not express any sulfatase 2 **(Figure 6E)**. We also performed pairwise comparisons using Dunn’s test and found there was a significant difference between the percentages of cells that were moderately positive for sulfatase 2 in tissue sections of TNBCs and ER+/PR+ breast cancers (p = 0.0497) **(Figure 6C)**. We found there was no significant difference when comparing the average percentages of cells that were strongly positive for sulfatase 2 between the different groups **(Figure 6D)**. We found that there was a significant difference between the percentages of cells that were positive for sulfatase 2 in tissue sections of TNBC breast cancer stages 2A vs. 2B (p = 0.0003) **(Figure 6F)**. Additional statistical analysis was carried out comparing the sulfatase 2 expression among normal breast tissue, DCIS, and various grades of cancers **(Supplemental Figure 11)**. We saw that high sulfatase 2 expression was associated with lack of expression of the PR **(Supplemental Figure 12)**. Furthermore, we observed that the higher percentage of Ki67 was associated with a higher expression of sulfatase 2 **(Supplemental Figure 13)**.

**Figure 6.**
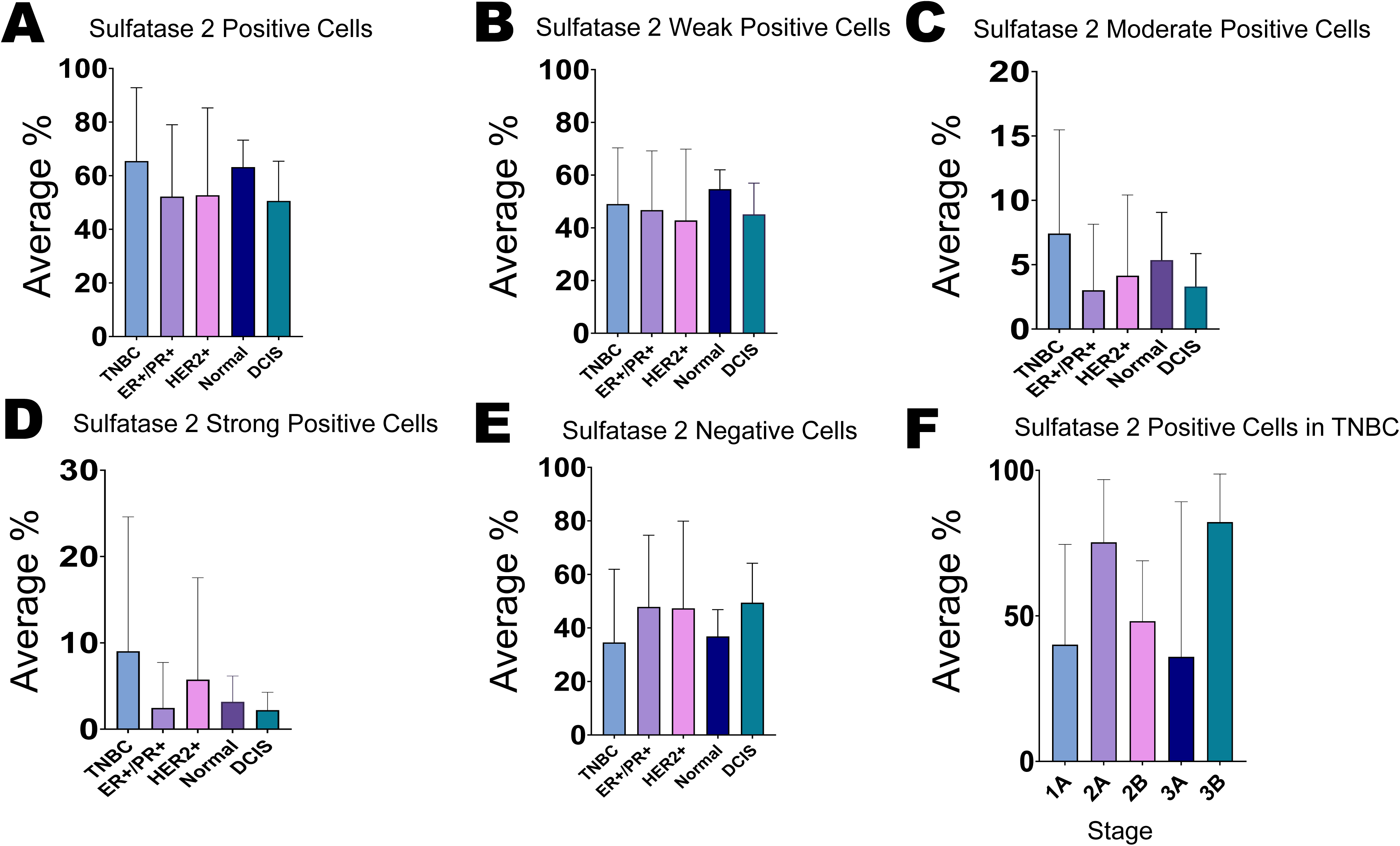
Statistical analysis for sulfatase 2 immunohistochemistry. (**A**) The Kruskal-Wallis test indicated that there was no significant difference in the average percentages of cells that stained positively for sulfatase 2 in tissue sections of TNBC, ER+/PR+ breast cancer, HER2+ breast cancer, normal breast tissue, and DCIS. **(B)** The Kruskal-Wallis test results showed that there was no significant difference between the average percentages of cells that stained weakly positive for sulfatase 2 in tissue sections of TNBC, ER+/PR+ breast cancer, HER2+ breast cancer, normal breast tissue, and DCIS. **(C**) Pairwise comparisons using Dunn’s test indicated that there was a significant difference between breast cancer sub-types TNBC and ER+/PR+ (*p* = 0.0497) in the percentages of cells that stained moderately positive for sulfatase 2. No other differences were statistically significantly between the other groups. **(D)** Pairwise comparisons using Dunn’s test showed that there was no significant difference between the percentages of cells that stained positive for sulfatase 2 among TNBC, ER+/PR+ breast cancer, HER2+ breast cancer, normal breast tissue, and DCIS. **(E)** The Kruskal-Wallis test indicated that there was no significant difference in the percentages of cells that stained negatively for sulfatase 2 among TNBC, ER+/PR+ breast cancer, HER2+ breast cancer, normal breast tissue, and DCIS. **(F)** Pairwise comparisons using Dunn’s test showed that there was a significant difference between TNBC breast cancer stages 2A and 2B (*p* = 0.0003) in the percentages of cells that stained positively for sulfatase 2. No other differences among different TNBC stages were significant.

**Figure 7.**
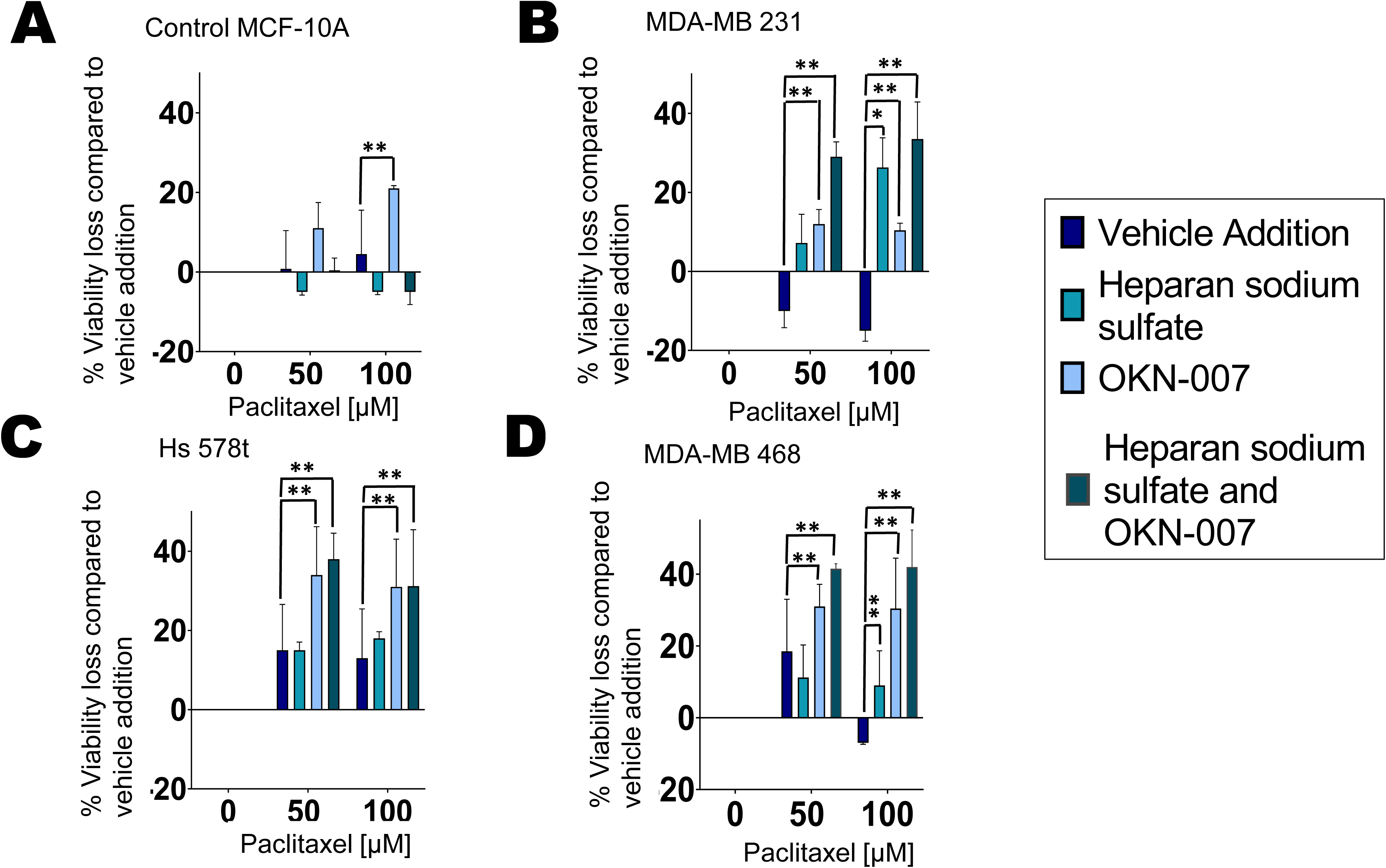
Effects of sulfatase inhibitor OKN-007 combined with chemotherapeutic agent paclitaxel on cell viability. Percentage loss of cell viability was measured in treated **(A)** nontumorigenic immortal mammary epithelial MCF-10A cells and TNBC cell lines **(B)** MDA-MB 231, **(C)** Hs 578t, and **(D)** MDA-MB 468 cells. The treatments applied were vehicle addition (paclitaxel, purple), heparan sodium sulfate (50 µM, teal), and OKN-007 (20 µM, light blue) or the combination (dark blue-green); heparan sodium sulfate and OKN-007 were administered for 48 hours, and paclitaxel was added for the final 6 hours to replicate paclitaxel exposure times in patients. Standard deviation was calculated from three independent experiments performed in triplicate. One-way ANOVA with Tukey’s HSD was applied to ascertain significance. * represents *p*<0.05 and ** represents *p*<0.01 when comparing vehicle addition to heparan sodium sulfate, OKN-007, or the combination.

### Effect of sulfatase inhibitor effect on cell viability and eATP

We previously demonstrated that eATP augmentation by ecto-ATPase inhibitors increases chemotherapy-induced TNBC cell death [8]. Given that polysulfated heparan is an ecto-ATPase inhibitor, we next determined the effect of combining the sulfatase inhibitor (OKN-007) with chemotherapy (paclitaxel) to ascertain its impact on the viability of TNBC MDA-MB 231, Hs 578t, and MDA-MB 468 cells in comparison to nontumorigenic immortal mammary epithelial MCF-10A cells and its effects on eATP release. For these experiments, all the cell lines were treated with OKN-007 and heparan sodium sulfate for 48 hours and then paclitaxel or a corresponding vehicle was added to the medium for the final 6 hours. The 6-hour duration of paclitaxel exposure was used to simulate the short duration of systemic exposure in cancer patients **(Figure 7)** [40]. As the treatment with paclitaxel was for only 6 hours, we did not see changes in the viability of cells treated with paclitaxel alone (vehicle addition). However, in all 3 TNBC cell lines—MDA-MB 231, Hs 578t, and MDA-MB 468—when paclitaxel (100 µM) was combined with the sulfatase inhibitor OKN-007 (20 µM), there was a significant loss of cell viability when compared to paclitaxel alone (shown as mean percentage loss of viability compared to vehicle control). Additionally, the loss of cell viability was further increased significantly when heparan sodium sulfate was added to the combination of paclitaxel and OKN-007 when compared to paclitaxel alone. However, there was no significant change in the viability of MCF-10A cells treated with paclitaxel, OKN-007, and heparan sodium sulfate compared to paclitaxel alone, suggesting that this effect may be selective for transformed cells **(Figure 7A)**.

Under the same conditions, we also measured the concentration of eATP in the supernatants of the chemotherapy-treated (paclitaxel) cells **(Figure 8)**. Upon treatment with the combination of OKN-007 and paclitaxel, we saw significant increases in eATP levels when compared to vehicle addition in MCF-10A cells as well as TNBC MDA-MB 231, Hs 578t, and MDA-MB 468. There was also a significant increase in eATP with the addition of heparan sodium sulfate to the combination of paclitaxel and OKN-007 when compared to the vehicle addition. Therefore, the sulfatase inhibitor OKN-007 significantly increased eATP release upon chemotherapy treatment and sensitized TNBC cell lines to chemotherapy treatment.

**Figure 8.**
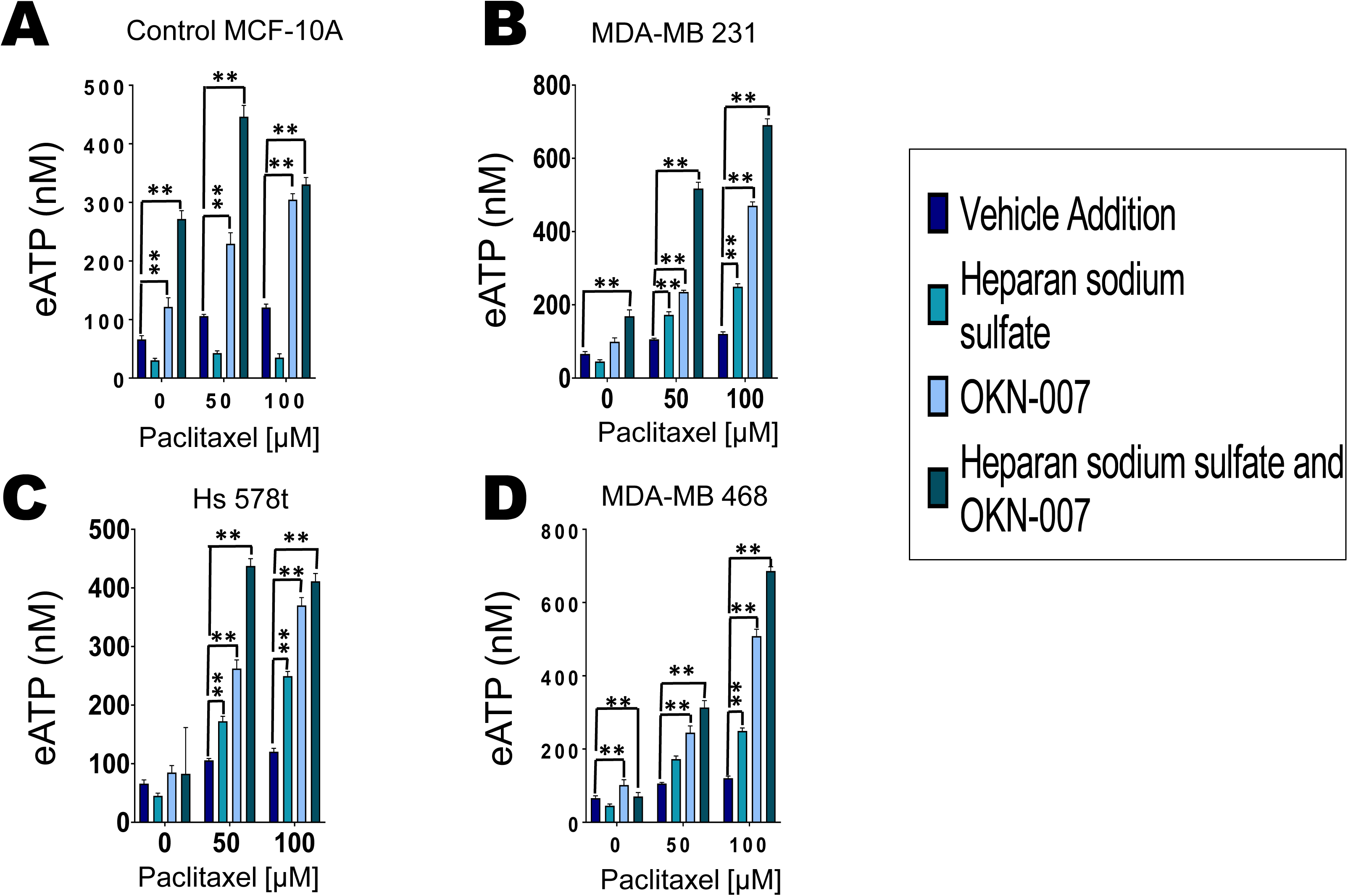
Effect of sulfatase inhibitor OKN-007, chemotherapeutic agent paclitaxel and their combination on extracellular ATP concentrations. Extracellular ATP concentrations were measured in the supernatants of treated **(A)** nontumorigenic immortal mammary epithelial MCF-10A cells and triple-negative breast cancer cell lines **(B)** MDA-MB 231**, (C)** Hs 578t, and **(D)** MDA-MB 468 cells. The treatments: vehicle addition (paclitaxel, purple), heparan sodium sulfate (50 µM, teal) and OKN-007 (20 µM, light blue), or the combination of all 3 (dark blue-green); heparan sodium sulfate and OKN-007 were administered for 48 hours and paclitaxel was added for the final 6 hours to replicate exposure times to paclitaxel in patients. Standard deviation was calculated from three independent experiments performed in triplicate. One-way ANOVA with Tukey’s HSD was applied to ascertain significance. * represents *p*<0.05 and ** represents *p*<0.01 when comparing vehicle addition to heparan sodium sulfate, OKN-007, or the combination regimen.

We also treated TNBC MDA-MB 231, Hs 578t and MDA-MB 468 cells and nontumorigenic immortal mammary epithelial MCF-10A cells paclitaxel (100 µM) for 6 hours or ATP (500 µM) for 48 hours and were probed for sulfatase 2 **(Supplemental Figures 14-16)**. We did not observe any change in sulfatase 2 expression in the presence of paclitaxel or ATP. We also obtained dose response graphs for treatments of increasing concentrations of paclitaxel and OKN-007 **(Supplemental Figures 17)**, There is some synergy (<0.1-1.0) for some dose combinations (paclitaxel and OKN-007) for MDA-MB 231 cells while there were some drug dose combinations (paclitaxel and OKN-007) that were additive (1-1.2) for Hs 578t and MDA-MB 468 cells.

### Role of purinergic signaling in the augmentation of chemotherapy-induced TNBC cell death by sulfatase 2 inhibitors

We had previously shown that eATP exerts its cytotoxic effects on TNBC cells through P2RX4 and P2RX7 receptors [8]. We sought to confirm whether the exaggerated loss of cell viability in the presence of OKN-007 is dependent on eATP-induced activation of P2RX4 or P2RX7 **(Figure 9)**. We chose to examine Hs 578t cells because we observed the largest increase in eATP and loss of cell viability in this cell type when exposed to the combination of paclitaxel, OKN-007, and heparan sodium sulfate. We observed a reversal of the effects of OKN-007 on cell viability and eATP release upon exposure to both the P2RX7 inhibitor A438079 **(Figures 8A and C)** and the P2RX4 inhibitor 5-BDBD **(Figures 9B and D)**. We observed a significant decline in eATP (p<0.0001) and increased cell viability (p<0.0001) when comparing the combination of paclitaxel with OKN-007 to that of paclitaxel with OKN-007 and A43709. We observed a significant decline in eATP (p<0.0001) and increased cell viability (p<0.0001) when comparing the combination of paclitaxel, heparan sodium sulfate and OKN-007 to paclitaxel, heparan sodium sulfate, OKN-007 and 5-BDBD. This data reveals that the exaggerated loss of cell viability observed when OKN-007 is combined with paclitaxel is dependent on the activation of P2RX4 and P2RX7 by eATP.

**Figure 9.**
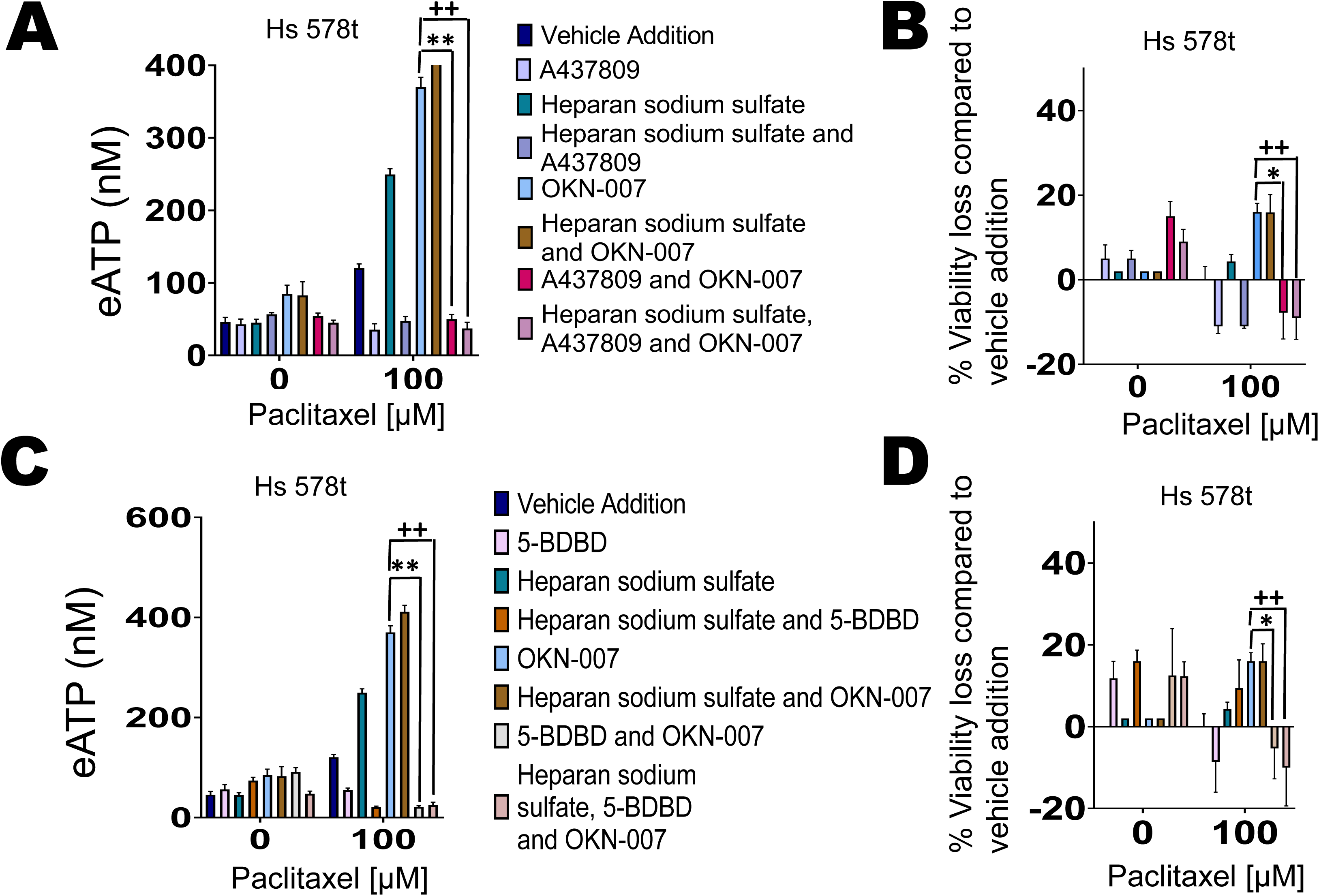
Reversal of sulfatase inhibitor’s effects by P2RX4 and P2RX7 inhibitors. For experiments corresponding to figures **(A)** and **(B),** Hs 578t cells were treated with OKN-007 (20 µM, 48 hours), paclitaxel (100 µM, the final 6 hours of the 48-hour time course to replicate exposure times in patients), heparan sodium sulfate (50 µM, 48 hours), the P2RX7 antagonist A437809 (20 µM, 6 hours), or a combination of the different drug agents. Standard deviation was calculated from three independent experiments performed in triplicate. One-way ANOVA with Tukey’s HSD was applied to ascertain significance. * represents *p*<0.05 and ** represents *p*<0.01 when comparing OKN-007 to the combination of vehicle addition, OKN-007, and A437809; + represents *p*<0.05 and ++ represents *p*<0.01 when comparing OKN-007 to the combination of vehicle addition, OKN-007, A437809, and heparan sodium sulfate. For experiments corresponding to figures **(C)** and **(D),** Hs 578t cells were treated with OKN-007 (20 µM, 48 hours), paclitaxel (100 µM, final 6 hours of the 48-hour time course to replicate exposure times in patients), heparan sodium sulfate (50 µM, 48 hours), the P2RX4 antagonist 5-BDBD (20 µM, 6 hours), or combinations. Standard deviation was calculated from three independent experiments performed in triplicate. One-way ANOVA with Tukey’s HSD was applied to ascertain significance. * represents *p*<0.05 and ** represents *p*<0.01 when comparing OKN-007 to the combination of vehicle addition, OKN-007, and 5-BDBD; + represents *p*<0.05 and ++ represents *p*<0.01 when comparing OKN-007 to the combination of vehicle addition, OKN-007, 5-BDBD, and heparan sodium sulfate.

### Purinergic signaling and cancer-initiating cells

Additionally, we sought to analyze the impact of the sulfatase inhibitor OKN-007 in combination with the chemotherapeutic agent paclitaxel on cancer-initiating cell properties of treated cells. Breast cancer-initiating cells have previously been shown to express aldehyde dehydrogenase (ALDH) intracellularly and CD44, but not CD24 at the cell surface [41–44]. Hence, flow cytometry analysis was performed on TNBC cell lines MDA-MB 231, MDA-MB 468, and Hs 578t treated with OKN-007 for 48 hours and paclitaxel (100 µM) for the final 6 hours to determine the fraction of residual cells with a cancer-initiating cell phenotype (cells that express high levels of ALDH and CD44 but do not express CD24). Paclitaxel alone increased the fraction of residual cancer-initiating cells while OKN-007 combined with paclitaxel depressed the fraction of cancer-initiating cells **(Figures 10A-C; Supplemental Figures 18-22)**.

**Figure 10.**
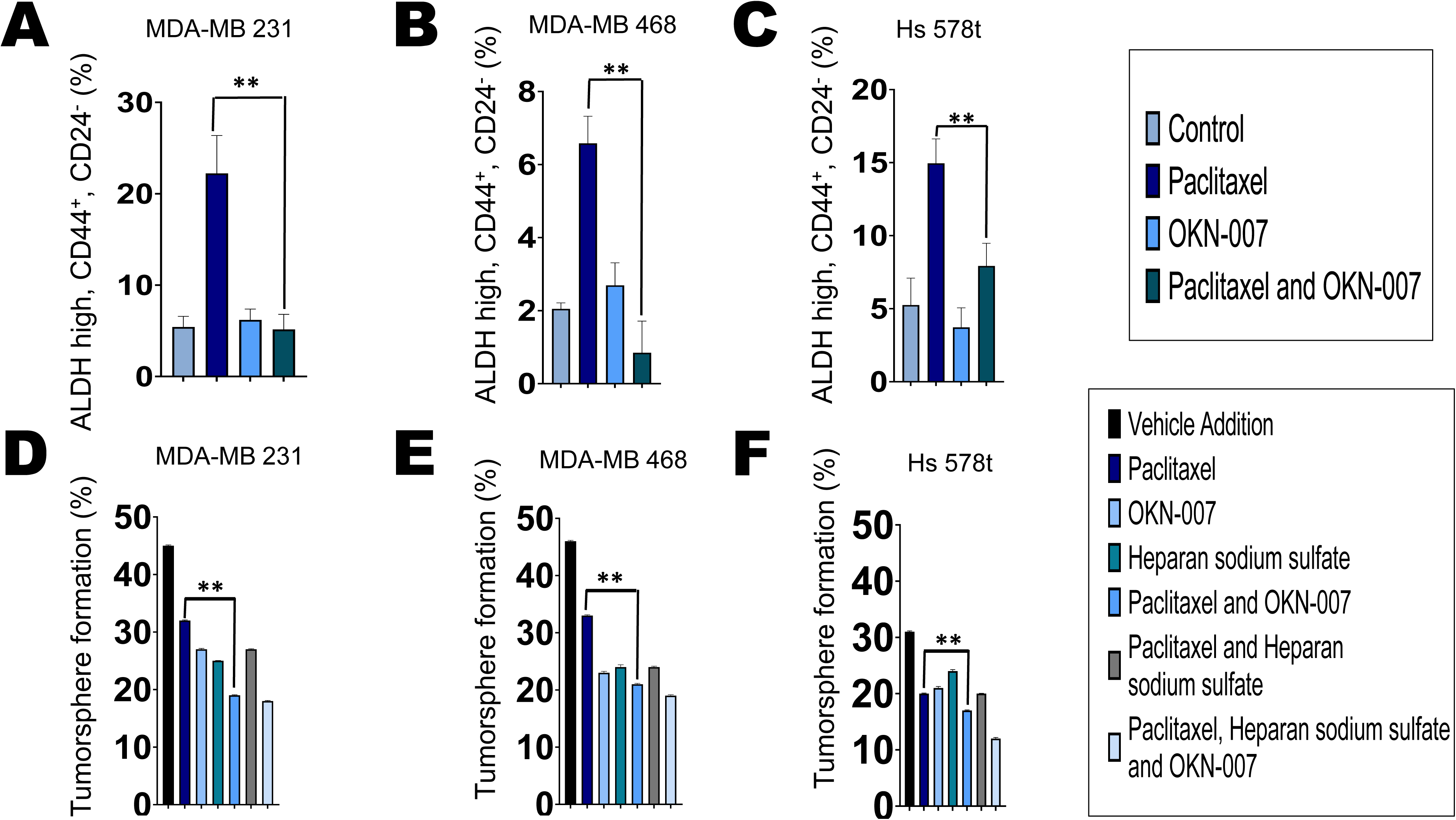
Effect of OKN-007 and chemotherapy on the cancer-initiating cell fraction of TNBC cells. The three TNBC cell lines **(A)** MDA-MB 231, **(B)** MDA-MB 468, and **(C)** Hs 578t were treated with OKN-007 (20 µM, 48 hours) and/or paclitaxel (100 µM, final 6 hours of the 48-hour time course to replicate exposure times in patients). Paclitaxel alone increased the cancer-initiating cell fraction among the surviving cells (cells that express high levels of ALDH and CD44 but do not express CD24), while the combination of OKN-007 and paclitaxel decreased the cancer-initiating cell fraction. One-way ANOVA with Tukey’s HSD was applied to ascertain significance. * represents *p*<0.05 and ** represents *p*<0.01 when comparing paclitaxel to paclitaxel and OKN-007. Effects on cancer-initiating cells were also determined through the tumorsphere formation efficiency assay in which TNBC cell lines **(D)** MDA-MB 231, **(E)** MDA-MB 468, and **(F)** Hs 578t cells were treated with vehicle (DMSO), paclitaxel (100 µM, final 6 hours of the 48-hour time course to replicate exposure times in patients), heparan sodium sulfate (50 µM, 48 hours), OKN-007 (20 µM, 48 hours), or the different combinations listed. Treated TNBC cells were washed, passed through cell strainers, collected, and plated at approximately one cell per well on round-bottom, low-attachment, 96-well plates; tumorspheres were allowed to form for 7 days. The fraction of wells plated with at least one live cell positive for tumorspheres after 7 days were counted using Etaluma™ Lumascope 620. The combination regimens showed a significant decrease in tumorsphere formation when compared to the single-agent treatments of vehicle, paclitaxel, heparan sodium sulfate, or OKN-007 treated cells. One-way ANOVA with Tukey’s HSD was applied to ascertain significance. ** represents *p*<0.01 when comparing paclitaxel to paclitaxel and OKN-007.

Furthermore, we applied an orthogonal approach to analyze cancer-initiating cells by conducting tumorsphere formation efficiency assays. TNBC cell lines were treated with paclitaxel, OKN-007, and/or heparan sodium sulfate and then cultured on low-attachment round-bottom plates in sphere-forming medium in the absence of drug. After 7 days, we observed that the fraction of wells with tumorsphere formation decreased in all TNBC cell lines—MDA-MB 231, MDA-MB 468, and Hs 578t—treated with paclitaxel as compared to vehicle control but was lowest when treated with the combination of paclitaxel and OKN-007. Spheroids were visualized using the Etaluma™ Lumascope 620 **(Figures 10D-F)**. Images were also recorded **(Supplemental Figures 23-25)**. This data aligns with flow cytometry analysis in that the sulfatase inhibitor OKN-007 depressed cancer-initiating cell formation.

## DISCUSSION

Chemotherapy is still the most effective treatment for TNBC. A major drawback of chemotherapy is its failure to eradicate metastatic disease, despite transient responses. Therapeutic strategies that deepen and lengthen responses are urgently needed. eATP, in the high micromolar to millimolar range, is cytotoxic to cancer cell lines. We previously showed that chemotherapy treatment augments eATP release from TNBC cells [8]. We also showed that ecto-ATPase inhibitors exacerbate chemotherapy-induced eATP release from TNBC cells and accentuate chemotherapy-induced cell death. However, one drawback of attempting to develop therapeutic small-molecule ecto-ATPase inhibitors is the presence of multiple families of ecto-ATPases in humans, each with multiple members.

Polysulfated polysaccharides have been shown to inhibit multiple classes of ecto-ATPases. The endogenous polysulfated HS was shown to attenuate the degradation of eATP. Therefore, we hypothesized that increasing fully sulfated HS in the microenvironment of TNBC cells using sulfatase 2 inhibitors would augment eATP concentrations in the pericellular environment of chemotherapy-treated TNBC cells, and hence accentuate chemotherapy-induced cell death.

Furthermore, our immunoblot results show that sulfatase 2 is highly expressed intracellularly and extracellularly in TNBC cells in comparison to the immortal mammary epithelial cells; this was additionally confirmed through ELISAs. However, the cell surface expression of sulfatase 2 did not differ among the TNBC cell lines and immortal mammary epithelial cells. Previous publications have suggested that processed sulfatase 2 is primarily an extracellular secreted protein [24, 26, 39]. This could explain the difference in the expression levels in cell supernatants as compared to cell surface expression. Staining results also revealed that there was a difference in the expression of sulfatase 2 at different stages of TNBC, with the most significant difference being between stages 2A and 2B.

Previously our lab showed that eATP in the high micromolar to millimolar range is toxic to TNBC cells and that inhibitors of each of the major classes of ecto-ATPases exacerbate chemotherapy-induced increases in eATP [8]. One defect of this approach as a potential therapeutic strategy is the need to use different inhibitors for each ecto-ATPase class to maximally augment eATP levels. As polysulfated polysaccharides such as HS inhibit all 3 families of ecto-ATPases, we sought to determine the effects of sulfatase 2 inhibitors, which block the desulfation of HS, on chemotherapy-induced augmentation of eATP and chemotherapy-induced TNBC cell death [8]. After verifying that the sulfatase 2 inhibitor OKN-007 inhibits sulfatase activity at the concentration utilized, we showed that the combination of OKN-007 and chemotherapy (paclitaxel) accentuated extracellular eATP concentrations and enhanced the chemotherapeutic response in TNBCs, leading to a greater loss of viability. Additionally, the effects of the sulfatase 2 inhibitor on eATP levels and TNBC cell death were reversed by specific inhibitors of P2RX4 and P2RX7 eATP receptors, confirming that these effects are dependent on these purinergic receptors; we have previously shown that these receptors are necessary for chemotherapy-induced eATP release from TNBC cells and its cytotoxic effects [8].

We also examined the effects of combinations of sulfatase 2 inhibitors and chemotherapy on cancer-initiating cells, as failure to eliminate these cells results in the failure of cytotoxic chemotherapy to eradicate metastatic TNBC [41, 43–45]. We found that upon treatment with chemotherapy alone, there was an enrichment of cancer-initiating cells, as assessed by flow cytometry, across all TNBC cell lines, while combination with the sulfatase inhibitor prevented their enrichment in the surviving fraction of cells.

As eATP is a known immune danger signal and its metabolite adenosine a potent immunosuppressant, further work is necessary to determine the immune effects of ecto-ATPase inhibition by HS using immunocompetent in vivo TNBC models. Moreover, further work is necessary to determine if the cytotoxic effects on TNBC cells occur through non-specific permeability of P2RX7 ion-coupled channels or through downstream activation of pyroptosis by P2RX7. Our future research will focus on this aspect of the effects of HS in the TME. In addition, the precise reasons for the different properties of sulfatase 1 and sulfatase 2, one a tumor suppressor in many biological contexts and the other an established oncogene, need to be determined. The basis for this may be complex and related to the precise identity of the cell surface and extracellular matrix proteins targeted by each enzyme. Thus, additional work is necessary in this area.

## CONCLUSION

Sulfatase 2 inhibition sensitizes TNBC cell lines to chemotherapy by enhancing eATP concentrations in the microenvironment of chemotherapy-treated cells. Combinations of sulfatase 2 inhibitors with chemotherapy may attenuate the cancer-initiating cell fraction, unlike chemotherapy alone. Thus, sulfatase 2 inhibitors may have the potential to induce deeper and more durable responses when combined with chemotherapy. Moreover, as eATP is a known immune danger signal, it will be critical to evaluate the immune effects of this strategy. Our future goals are to study the effects of sulfatase 2 inhibition in vivo.

## DECLARATIONS

### Ethics approval and consent to participate

Not applicable.

### Consent for publication

Not applicable.

### Availability of data and materials

The datasets used and/or analyzed during the current study are available from the corresponding author upon reasonable request.

## Funding

Research reported in this publication was supported by The Ohio State University Comprehensive Cancer Center and by a National Cancer Institute (NCI)/National Institutes of Health (NIH) grant (P30CA016058). This publication was also supported, in part, by a grant from the National Center for Advancing Translational Sciences of the NIH (KL2TR002734). Institutions that provided funding support had no role in the design or conduct of this study or the preparation of the manuscript. The content is solely the responsibility of the authors and does not necessarily represent the official views of the NIH.

### Authors’ contributions

All authors contributed to review and analysis. JM performed a majority of the assays with JD carrying out the Western blot analysis, and LM carrying out immunohistochemistry analysis. JM and MC conceived of and designed the experiments, reviewed the data, and authored and edited the manuscript.

## ABBREVIATIONS

ALDH: aldehyde dehydrogenase
ANOVA: analysis of variance
ATP: adenosine triphosphate
DCIS: ductal carcinoma in situ
DMSO: dimethyl sulfoxide
eATP: extracellular adenosine triphosphate
ELISA: enzyme-linked immunoassay
E-NPPase: ecto-nucleotide pyrophosphatases/phosphodiesterases
ENPP1: ecto-nucleotide pyrophosphatase/phosphodiesterase 1
ENTPD1: ecto-nucleoside triphosphate diphosphohydrolases 1
ER+: estrogen receptor-positive
FACS: fluorescence-activated cell sorting
FITC: fluorescein isothiocyanate
5’-NTs: 5’-nucleotidases
GAPDH: glyceraldehyde-3-phosphate dehydrogenase
HIER: heat-induced epitope retrieval
HER2+: human epidermal growth factor receptor 2-positive
HS: heparan sulfate
HSD: honestly significant difference
MFI: mean fluorescence intensity
mM: millimolar
nM: nanomolar
PAGE: polyacrylamide gel electrophoresis
PBS: phosphate buffered saline
PR+: progesterone receptor-positive
TMA: tissue microarray
TME: tumor microenvironment
TNAP: tissue nonspecific alkaline phosphatases
TNBC: triple-negative breast cancer

## Acknowledgments

The authors would like to thank scientific editor Angela Dahlberg, Division of Medical Oncology, The Ohio State University Comprehensive Cancer Center, for editing this manuscript.

