## Supplemental Figures for "Sulfatase 2 Inhibition Sensitizes Triple-Negative Breast Cancer Cells to Chemotherapy Through Augmentation of Extracellular ATP"

### Slide 1
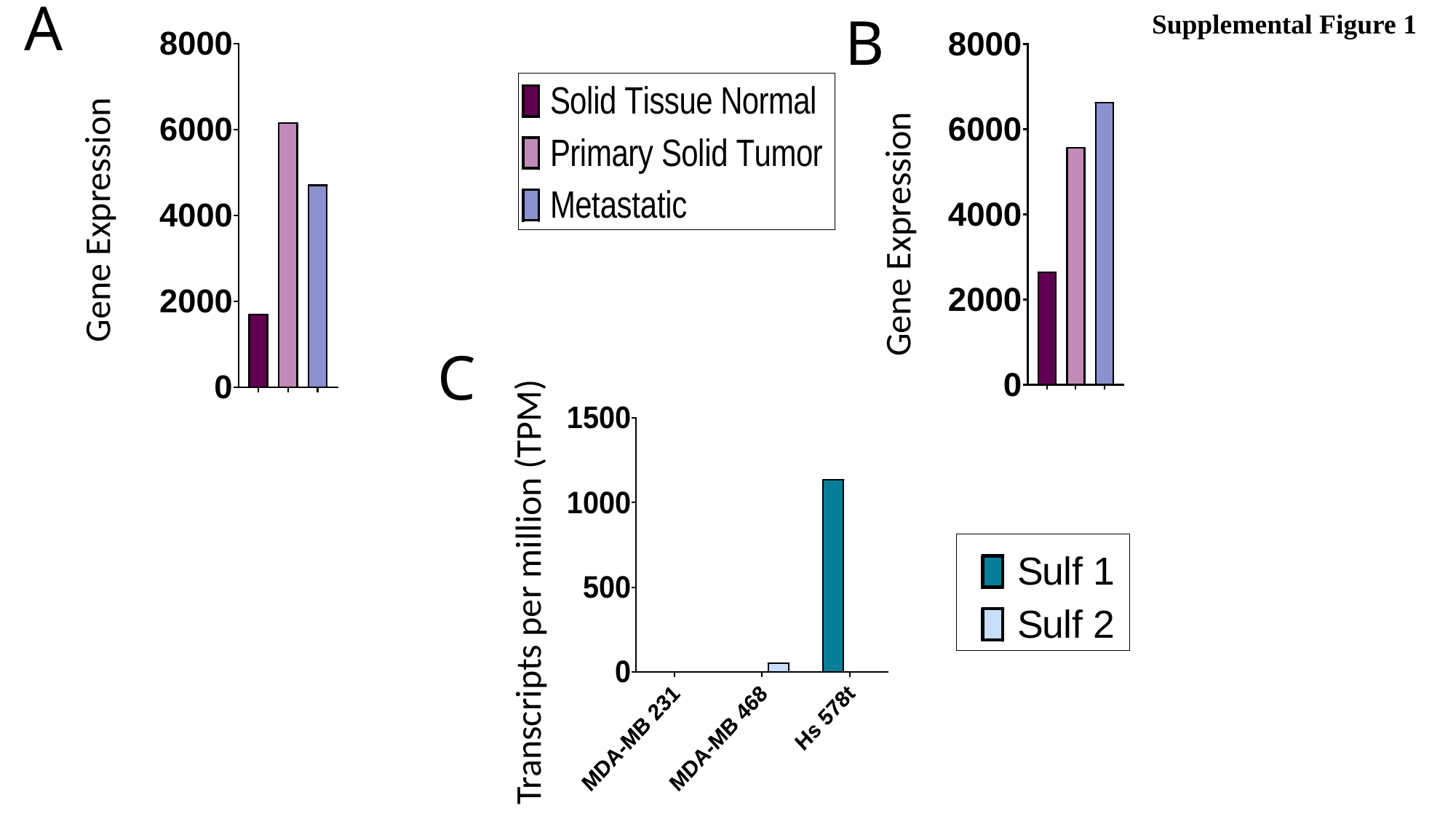

A
B
Supplemental Figure 1
Gene Expression
Gene Expression
C
Transcripts per million (TPM)

### Slide 2
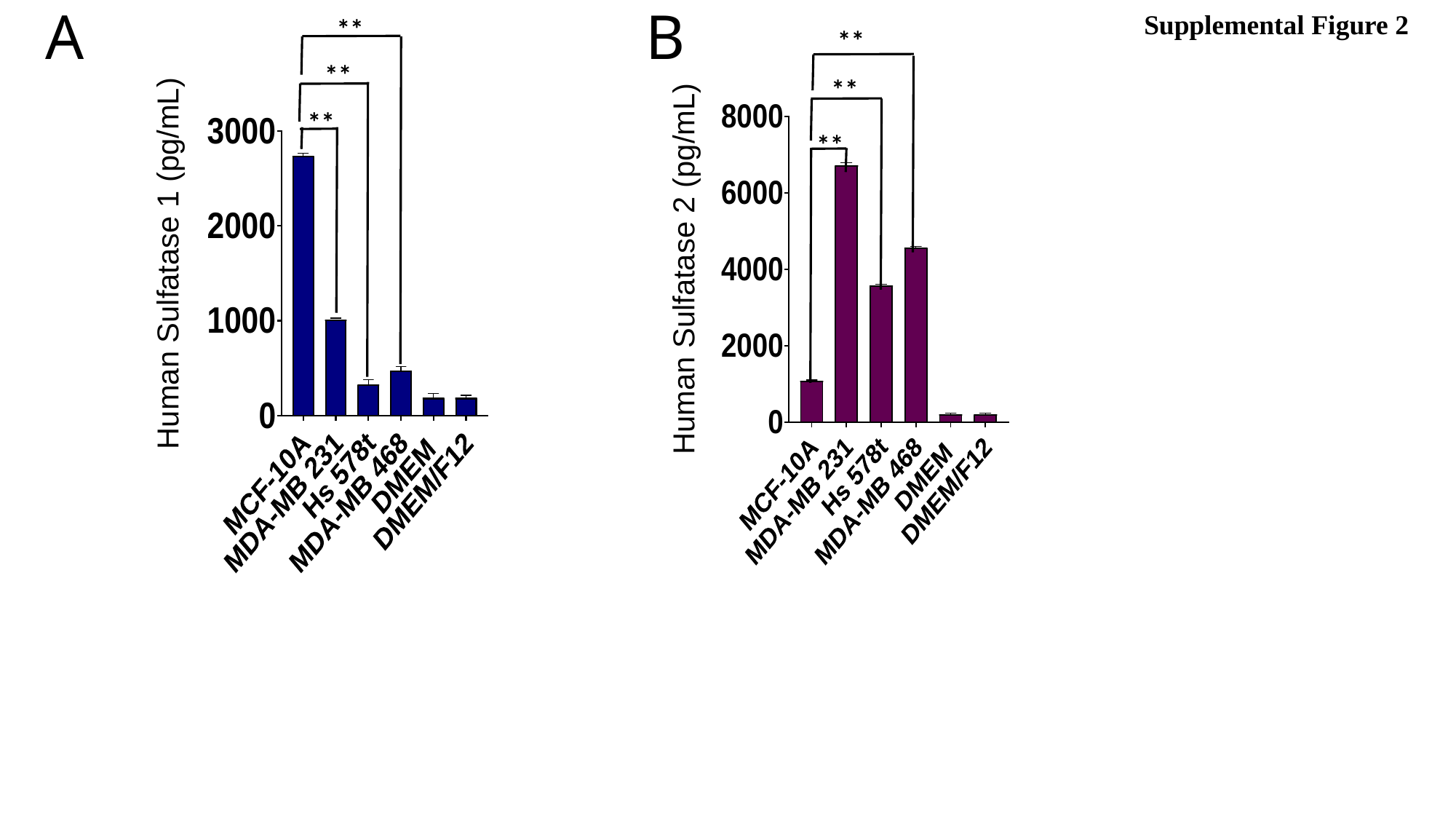

B
A
Supplemental Figure 2
**
**
Human Sulfatase 1 (pg/mL)
**
Human Sulfatase 2 (pg/mL)
**
**
**

### Slide 3
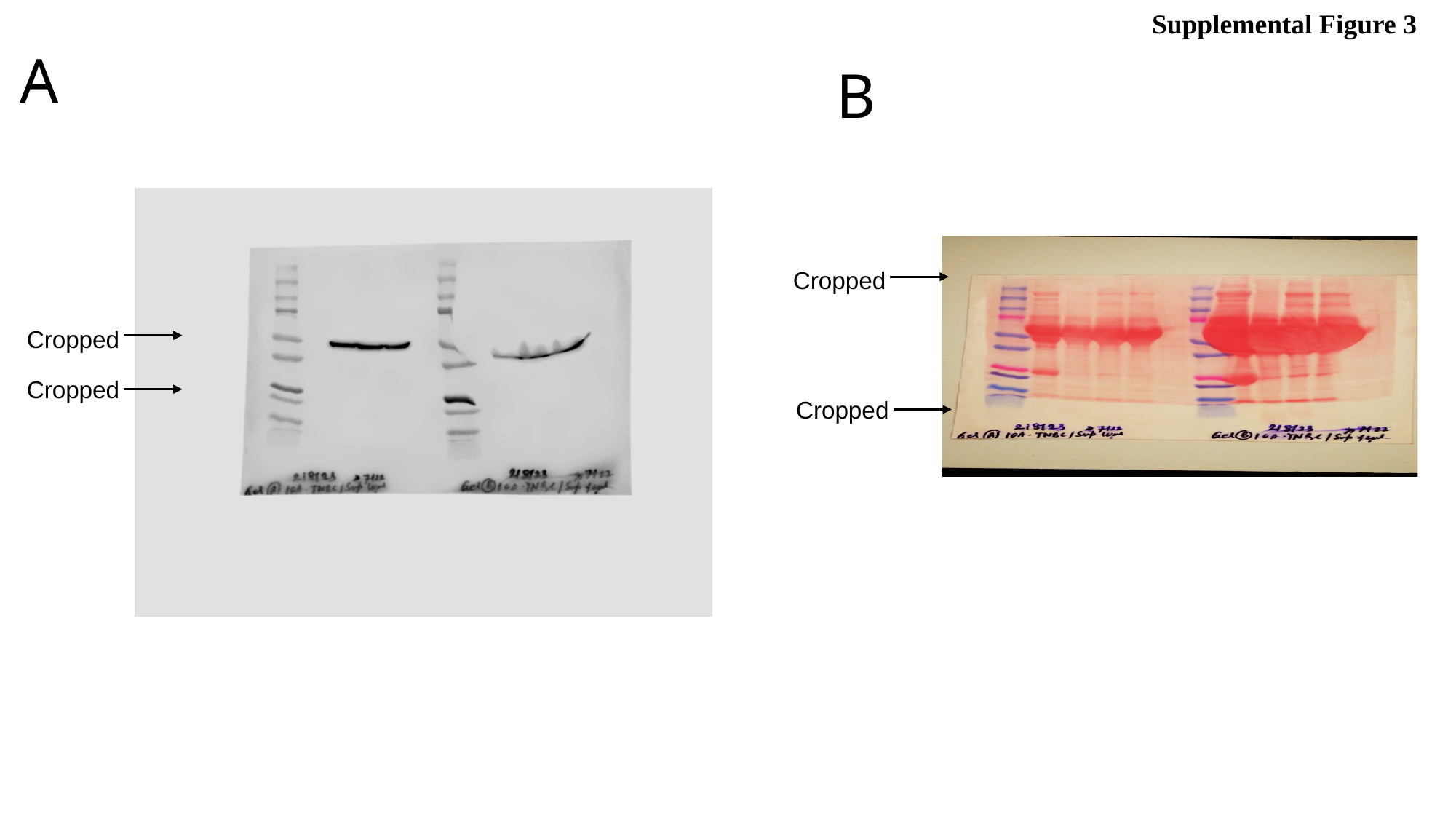

Supplemental Figure 3
A
B
 Cropped
 Cropped
 Cropped
 Cropped

### Slide 4
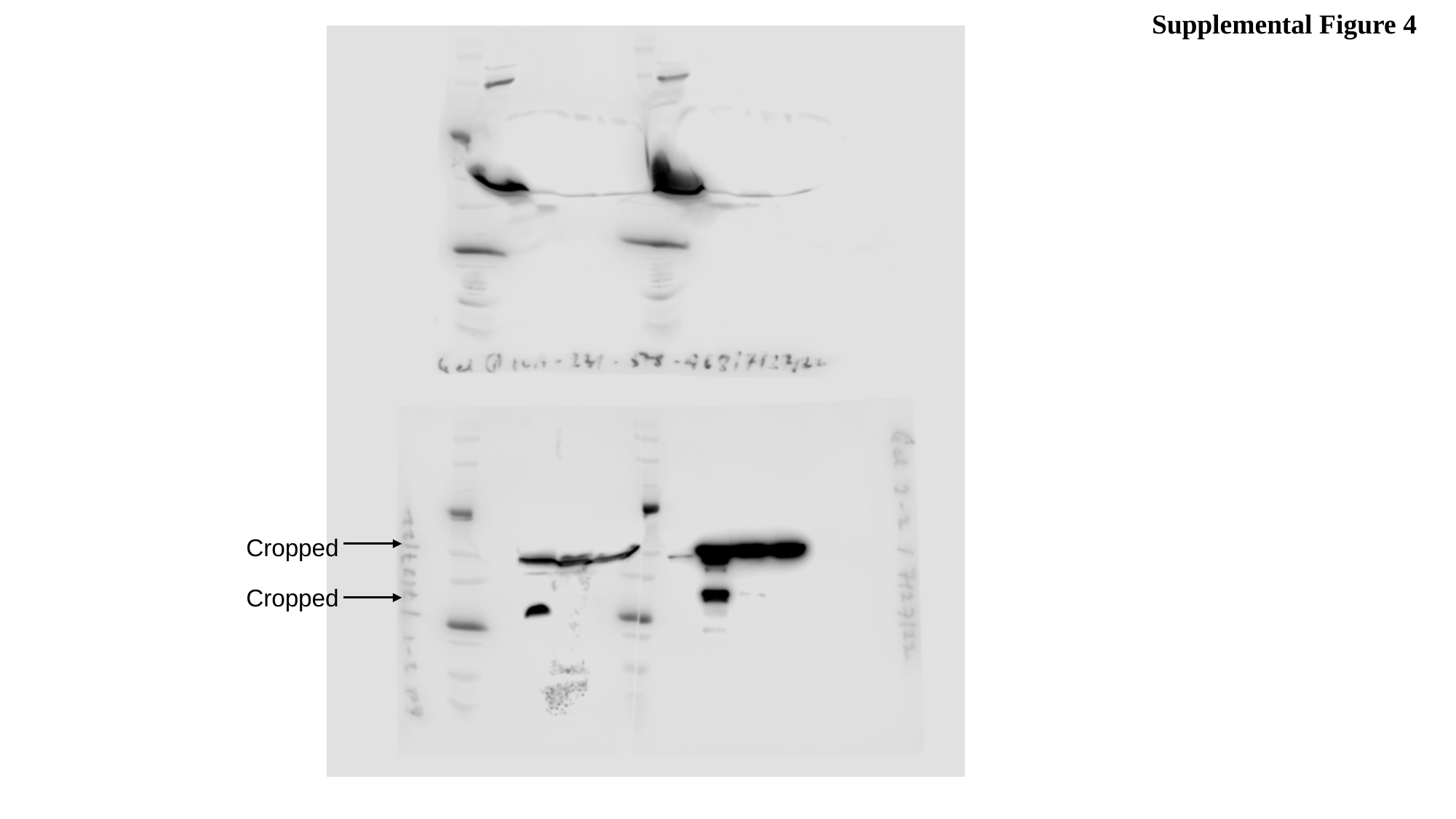

Supplemental Figure 4
 Cropped
 Cropped

### Slide 5
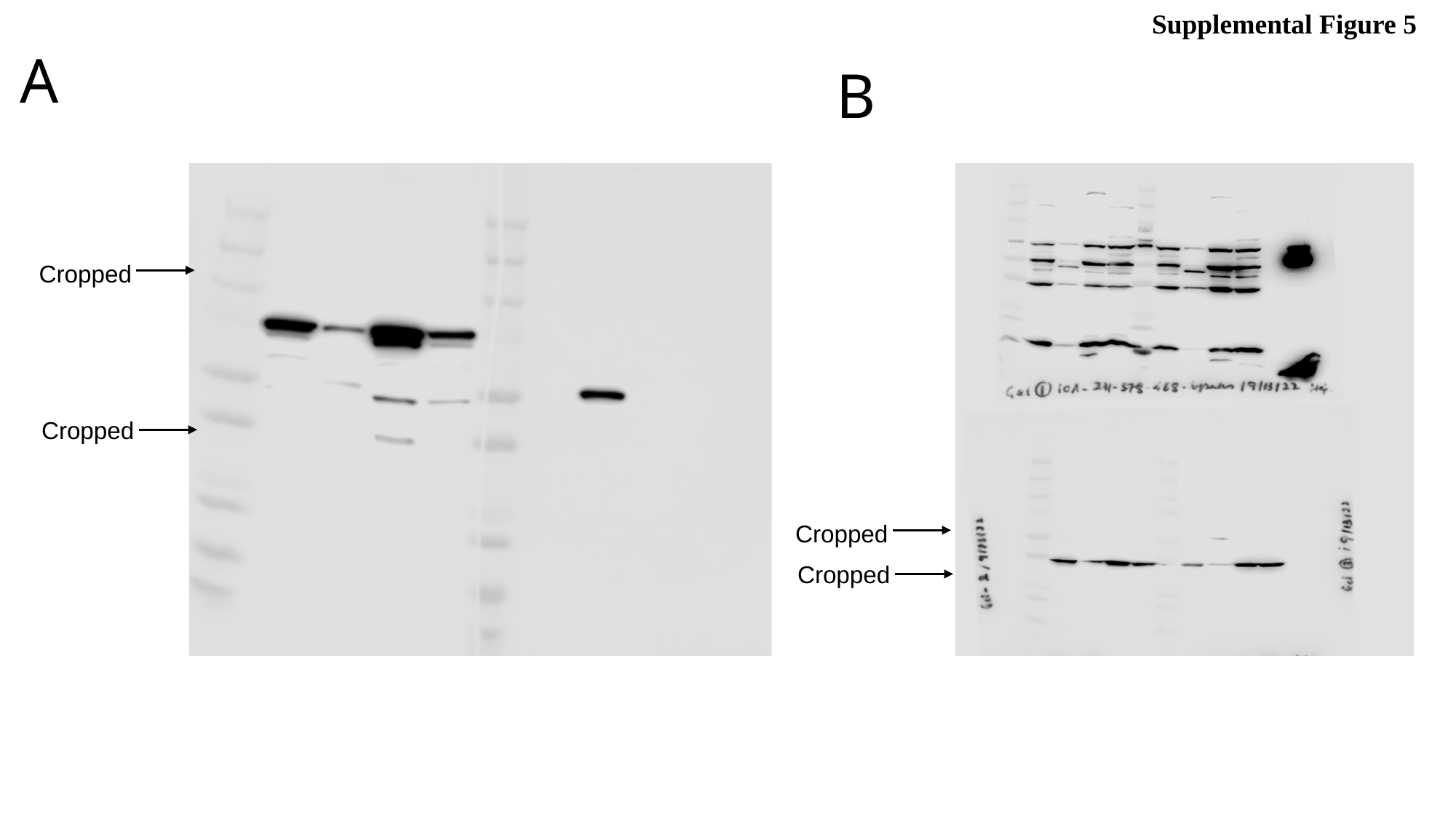

Supplemental Figure 5
A
B
 Cropped
 Cropped
 Cropped
 Cropped

### Slide 6
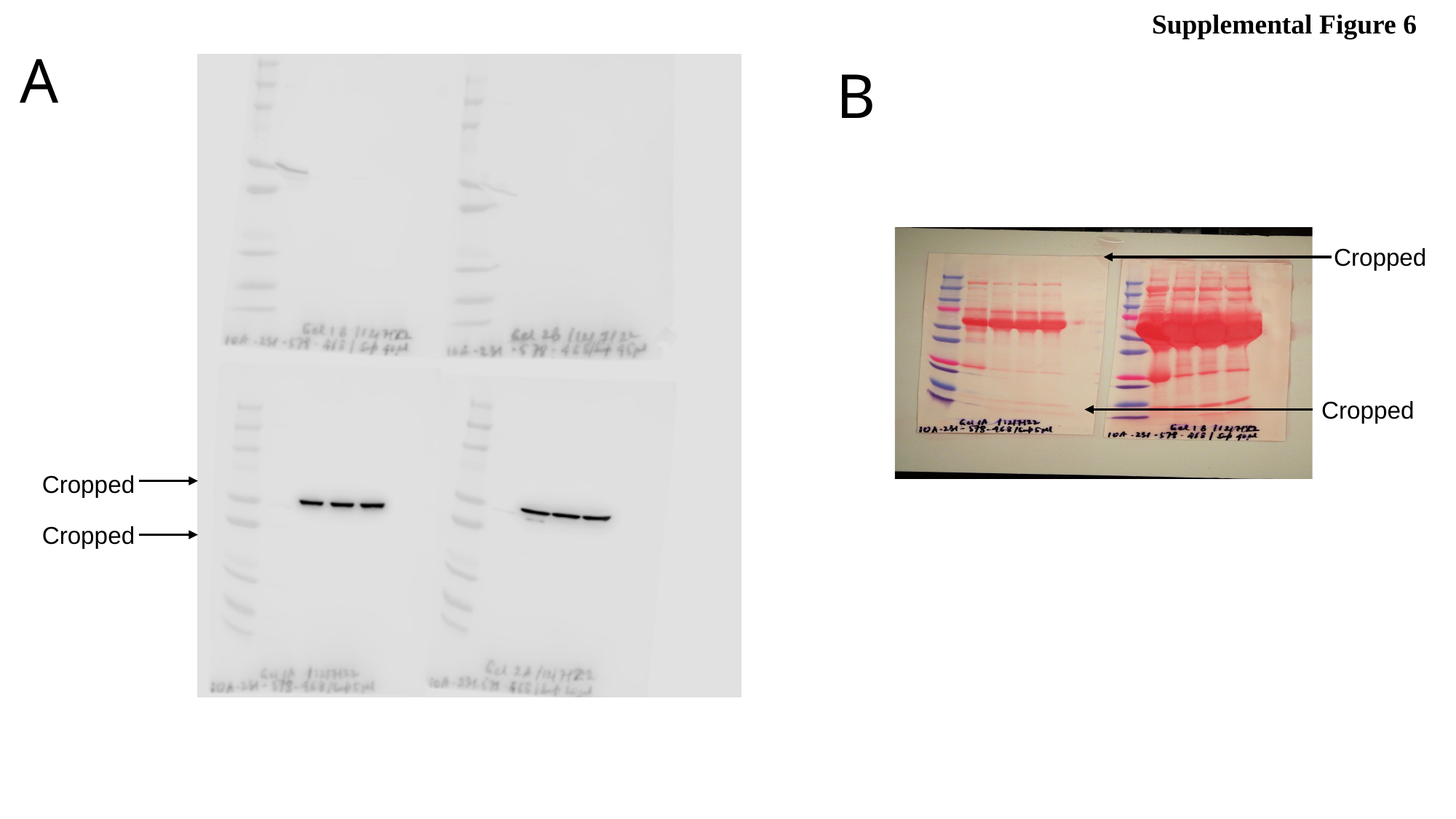

Supplemental Figure 6
A
B
 Cropped
 Cropped
 Cropped
 Cropped

### Slide 7
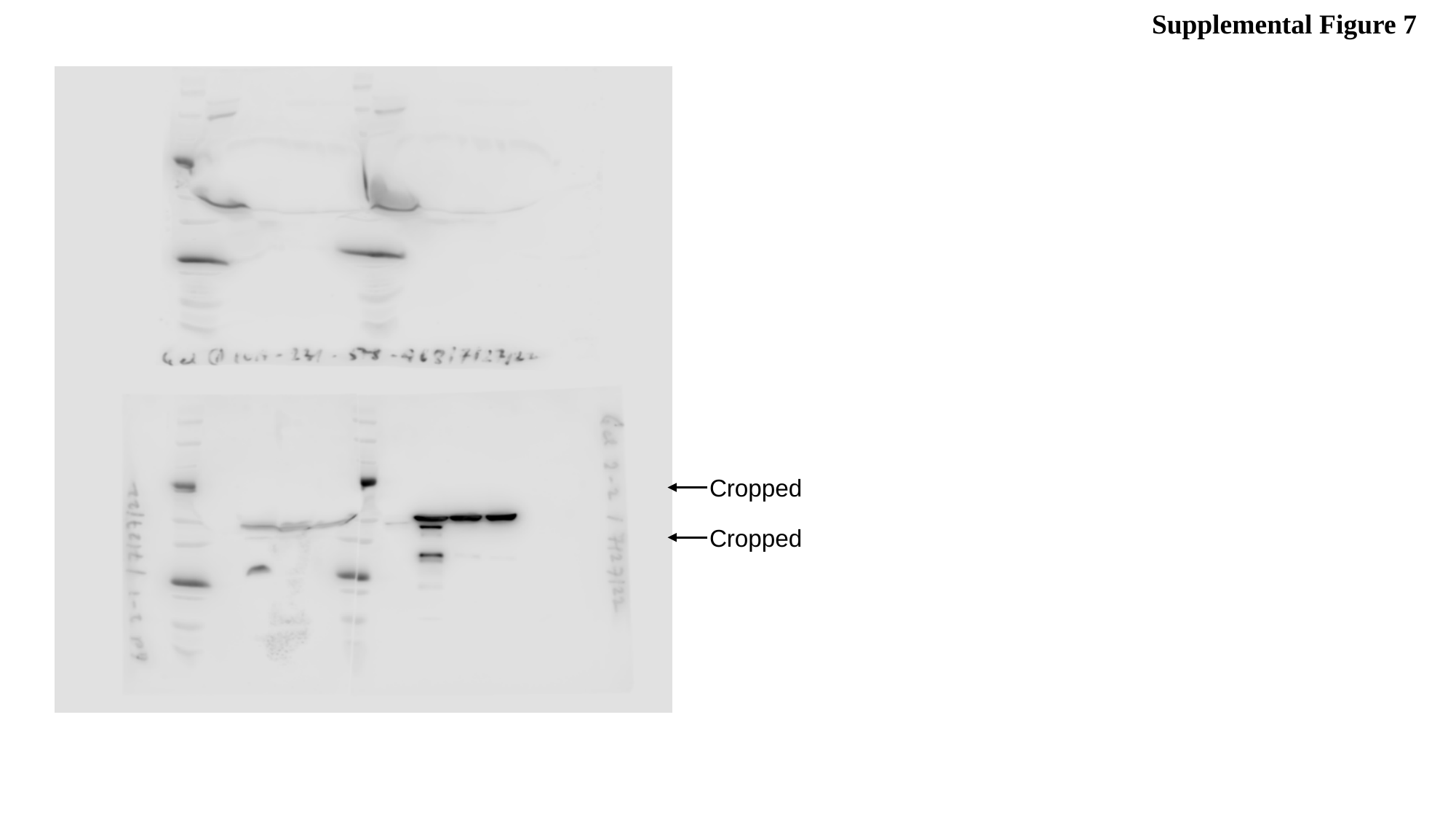

Supplemental Figure 7
 Cropped
 Cropped

### Slide 8
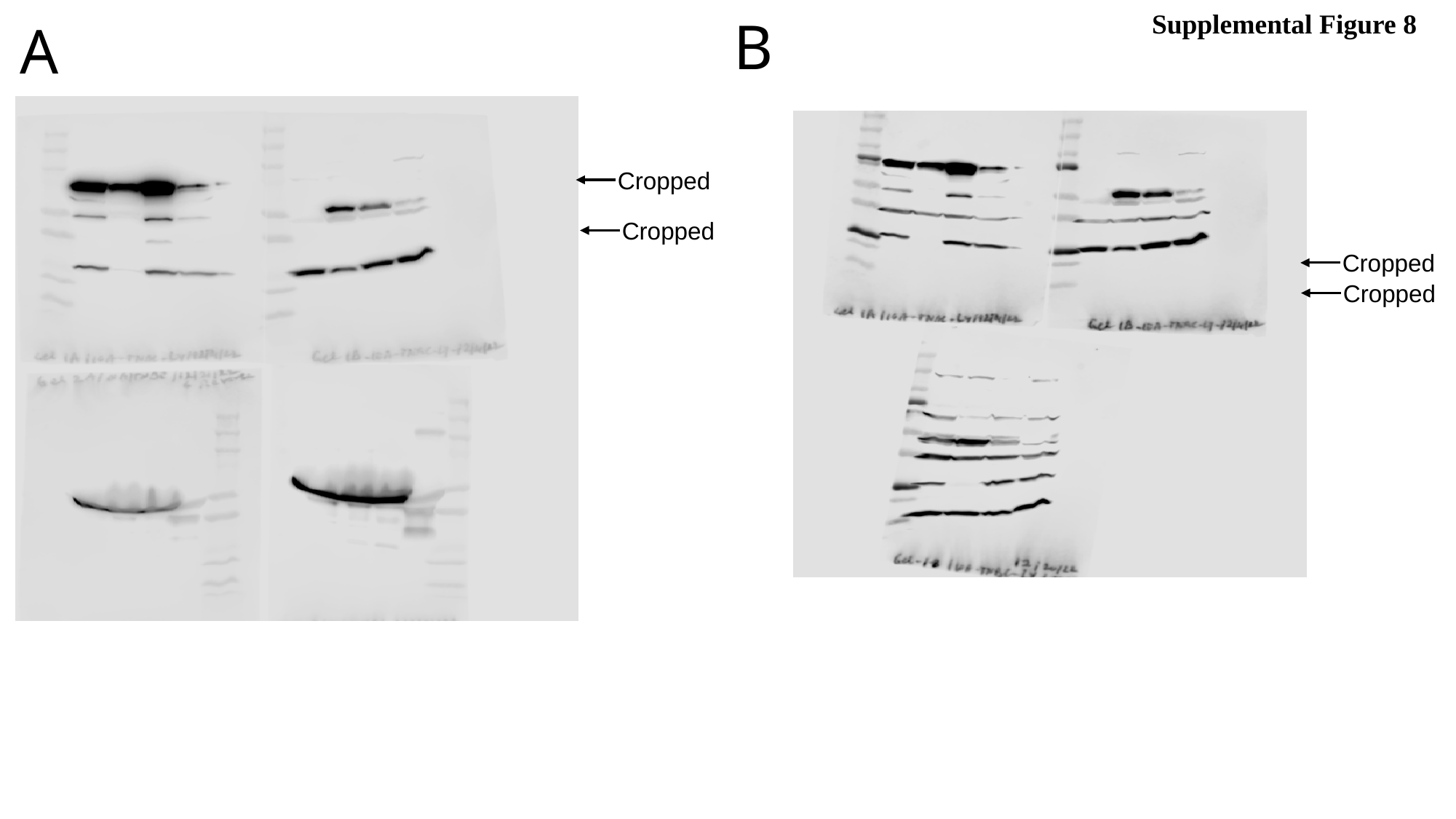

Supplemental Figure 8
B
A
 Cropped
 Cropped
 Cropped
 Cropped

### Slide 9
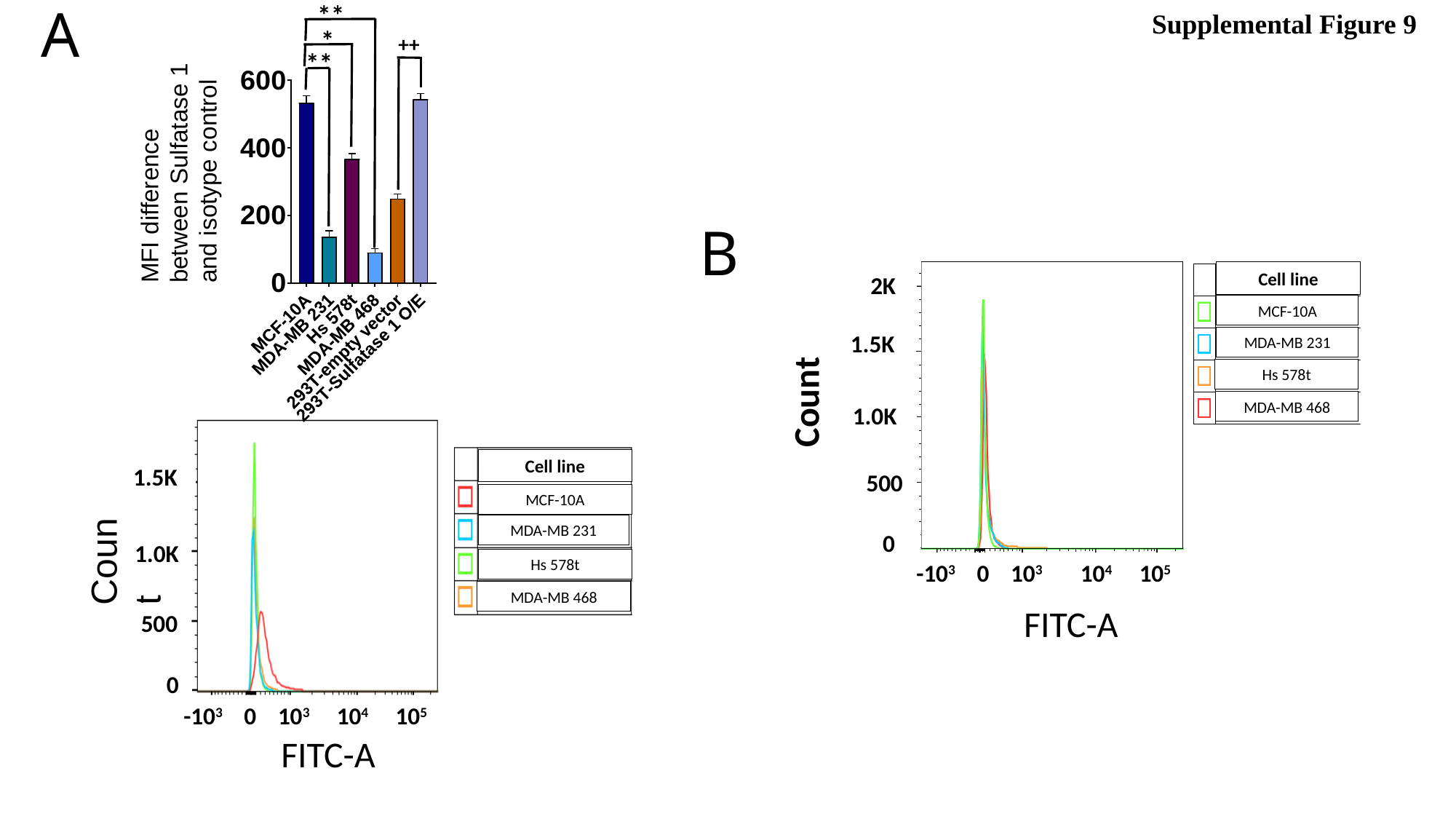

A
**
Supplemental Figure 9
*
MFI difference between Sulfatase 1 and isotype control
++
**
B
Cell line
2K
MCF-10A
Count
1.5K
MDA-MB 231
Hs 578t
MDA-MB 468
1.0K
Cell line
1.5K
500
MCF-10A
Count
MDA-MB 231
0
1.0K
Hs 578t
-10^3^ 0 10^3^ 10^4^ 10^5^
MDA-MB 468
FITC-A
500
0
-10^3^ 0 10^3^ 10^4^ 10^5^
FITC-A

### Slide 10
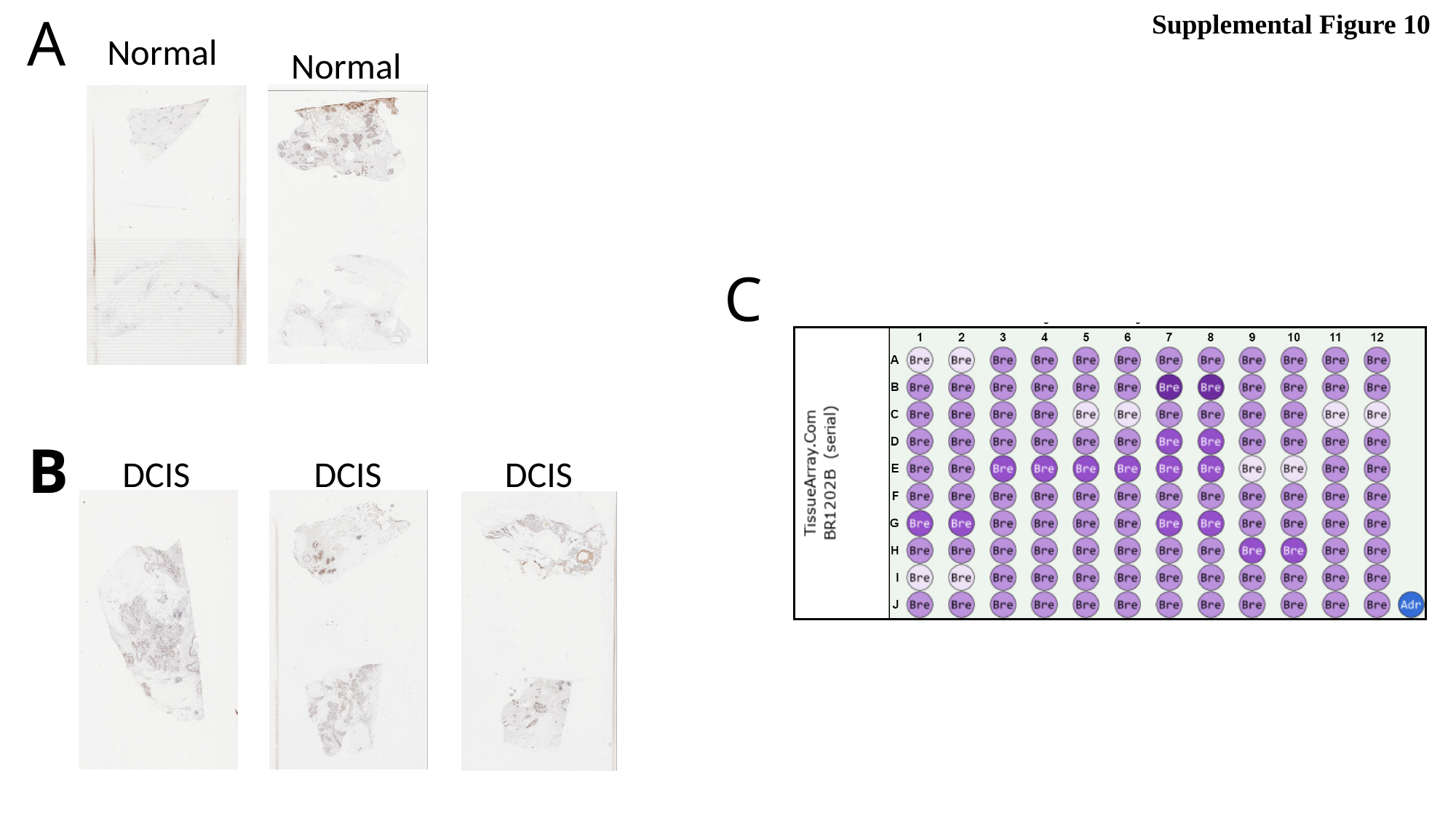

A
Supplemental Figure 10
Normal
Normal
C
B
DCIS
DCIS
DCIS

### Slide 11
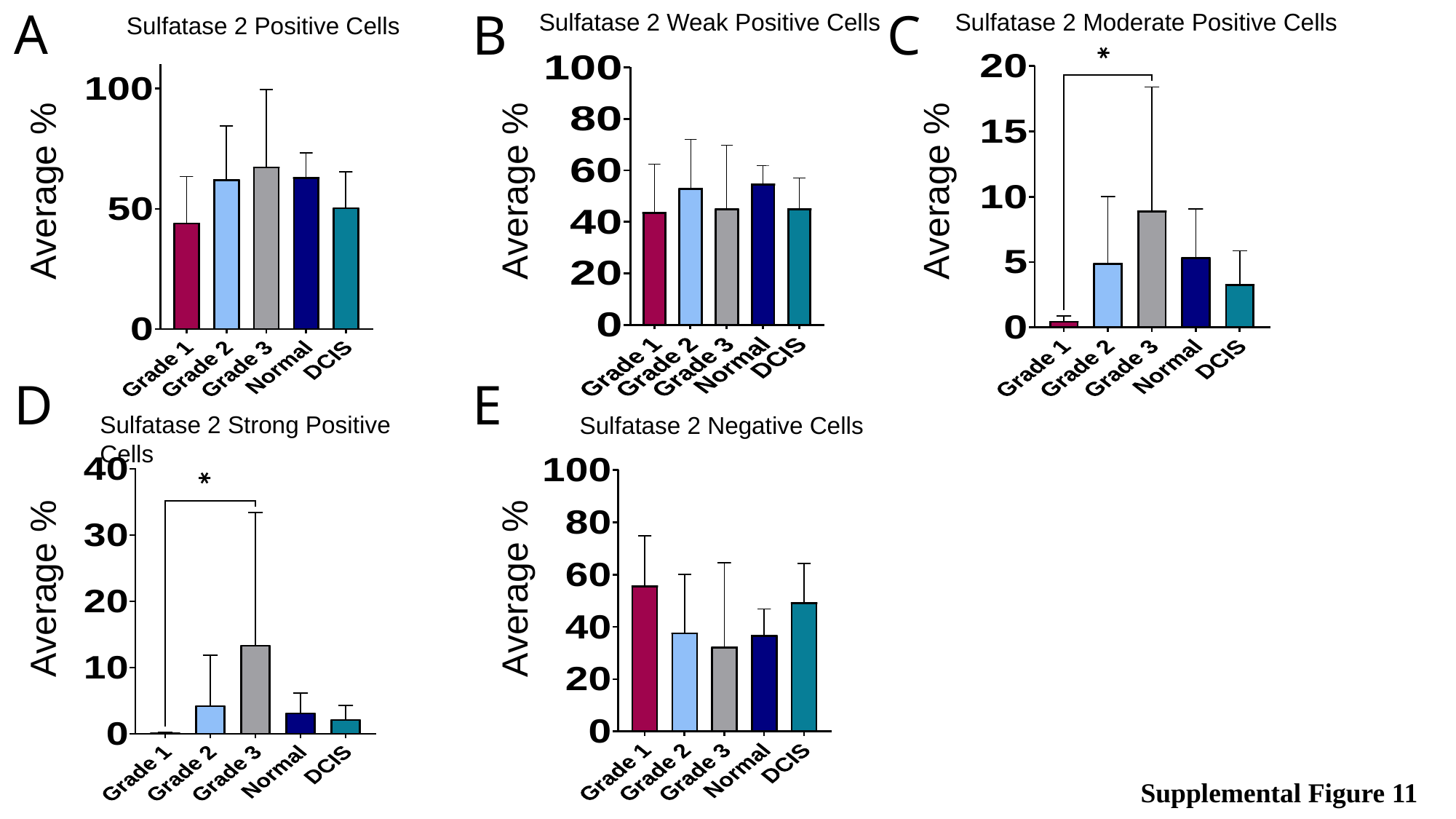

Sulfatase 2 Moderate Positive Cells
Sulfatase 2 Positive Cells
Sulfatase 2 Weak Positive Cells
A
B
C
*
Average %
Average %
Average %
Sulfatase 2 Strong Positive Cells
Sulfatase 2 Negative Cells
E
D
*
Average %
Average %
Supplemental Figure 11

### Slide 12
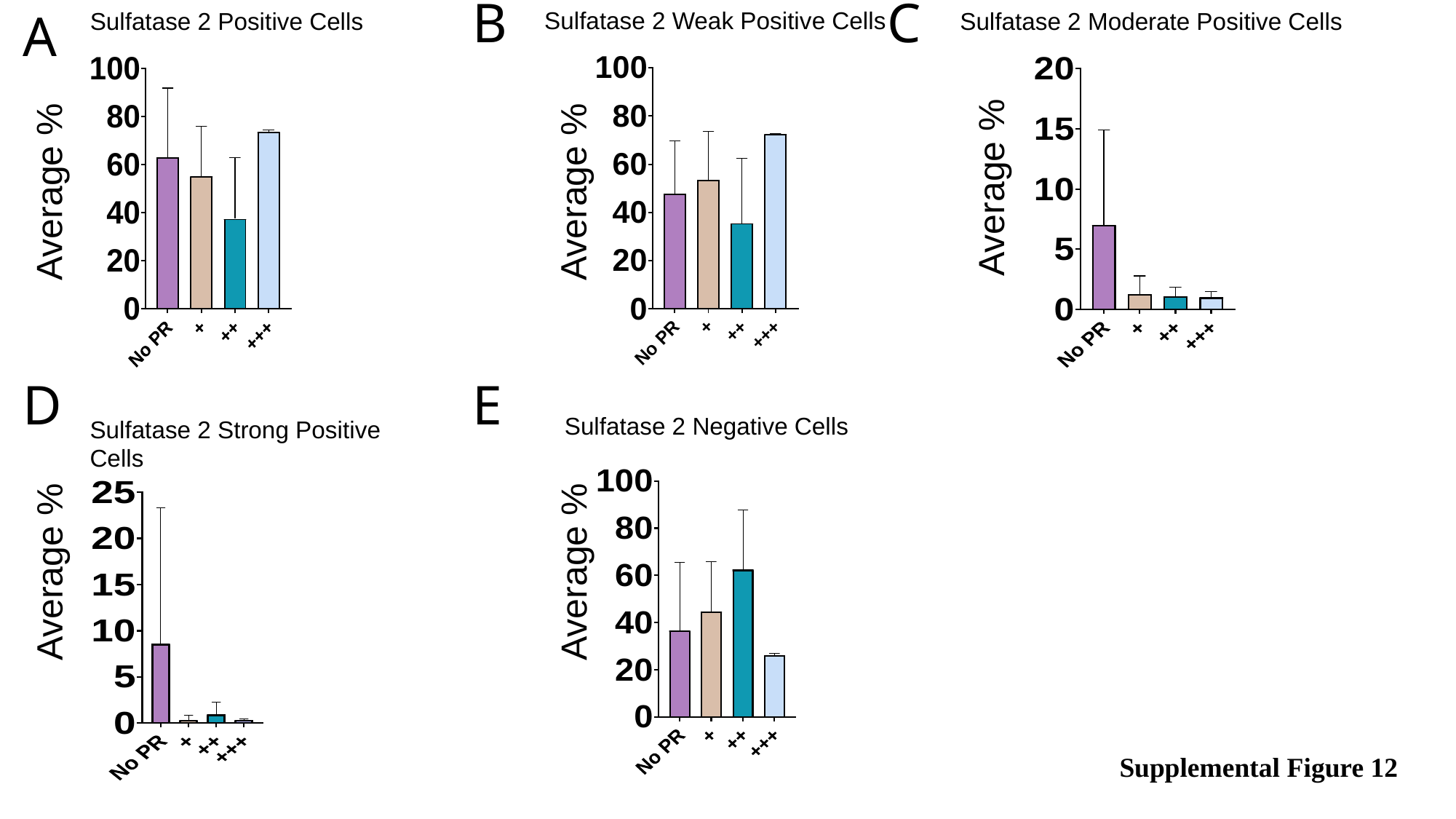

Sulfatase 2 Moderate Positive Cells
Sulfatase 2 Weak Positive Cells
Sulfatase 2 Positive Cells
B
C
A
Average %
Average %
Average %
Sulfatase 2 Strong Positive Cells
Sulfatase 2 Negative Cells
E
D
Average %
Average %
Supplemental Figure 12

### Slide 13
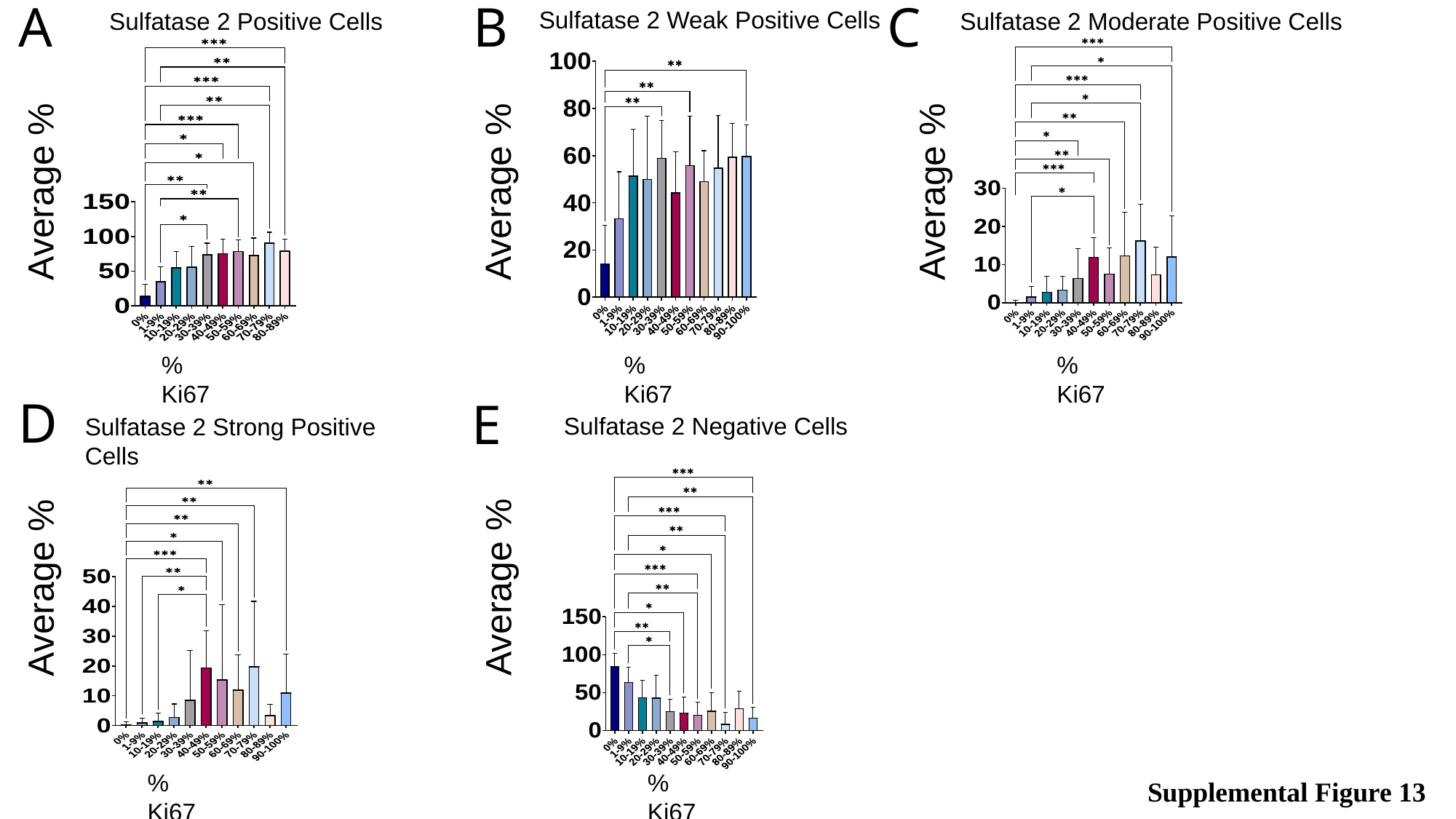

Sulfatase 2 Moderate Positive Cells
Sulfatase 2 Weak Positive Cells
Sulfatase 2 Positive Cells
B
C
A
Average %
Average %
Average %
Sulfatase 2 Strong Positive Cells
Sulfatase 2 Negative Cells
% Ki67
% Ki67
% Ki67
D
E
Average %
Average %
% Ki67
% Ki67
Supplemental Figure 13

### Slide 14
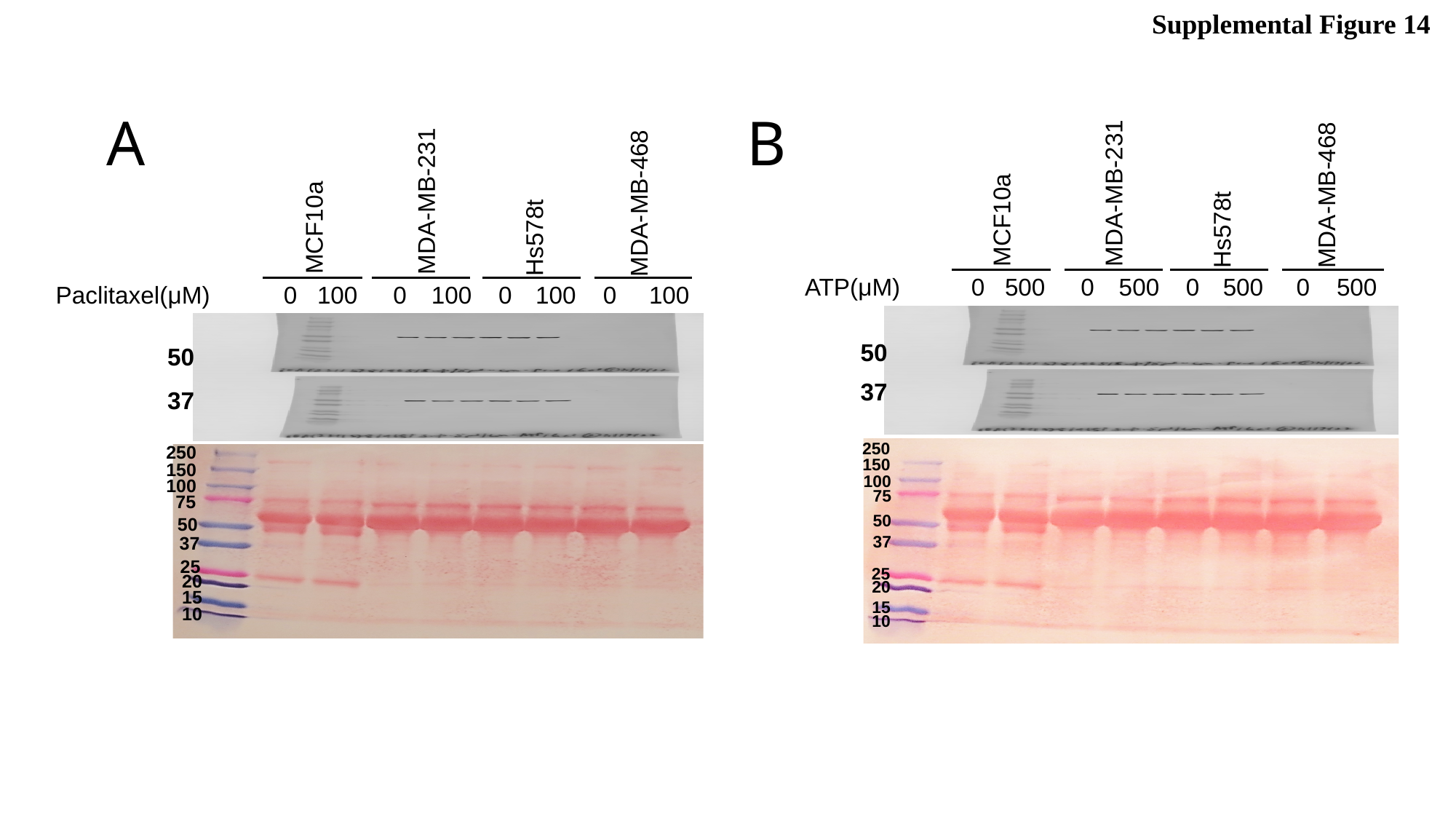

Supplemental Figure 14
A
MDA-MB-231
Hs578t
MDA-MB-468
MCF10a
0
100
0
100
0
100
0
100
Paclitaxel(μM)
50
37
250
150
100
75
50
37
25
20
15
10
B
MDA-MB-231
Hs578t
MDA-MB-468
MCF10a
0
500
0
500
0
500
0
500
ATP(μM)
50
37
250
150
100
75
50
37
25
20
15
10

### Slide 15
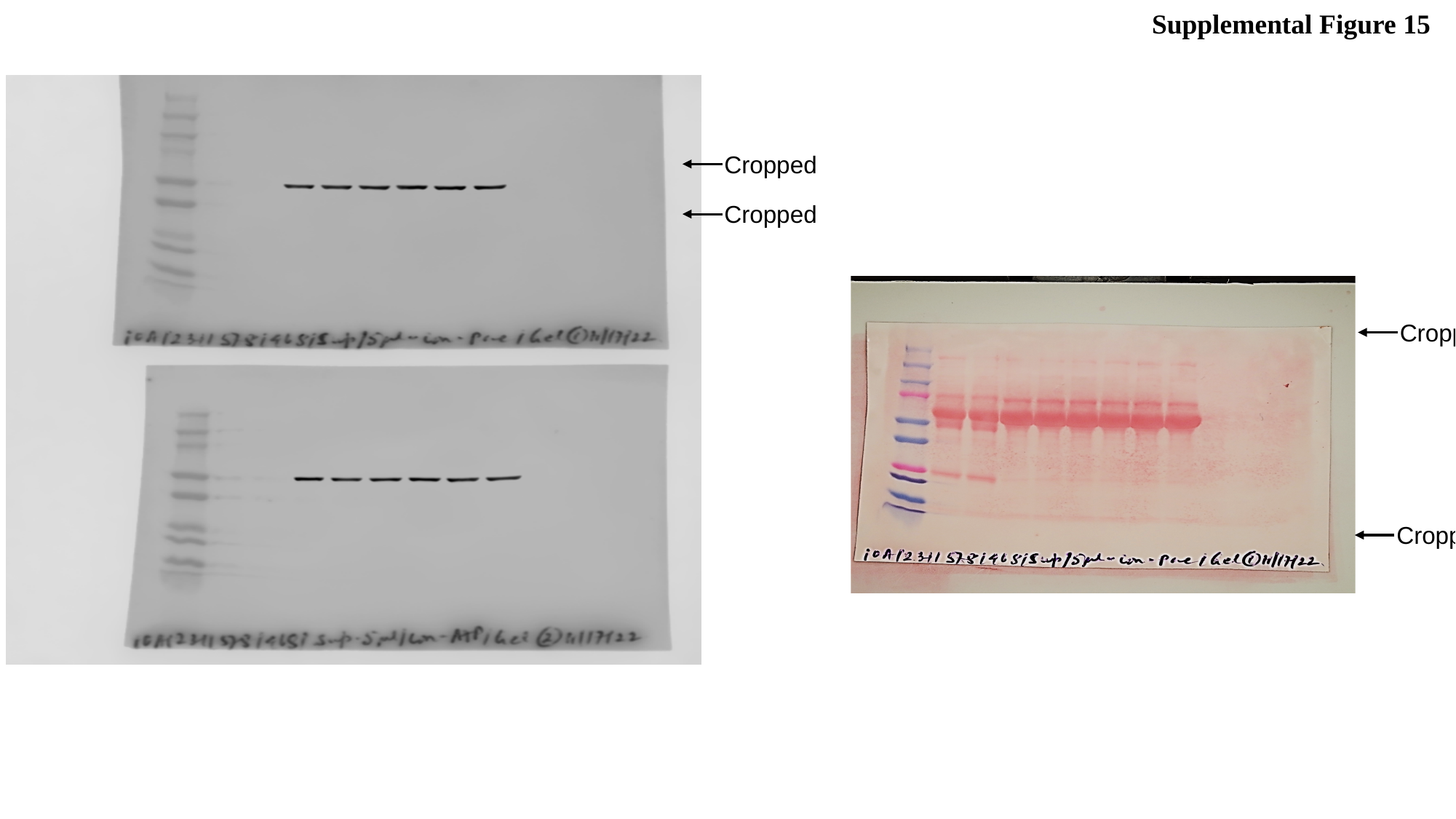

Supplemental Figure 15
 Cropped
 Cropped
 Cropped
 Cropped

### Slide 16
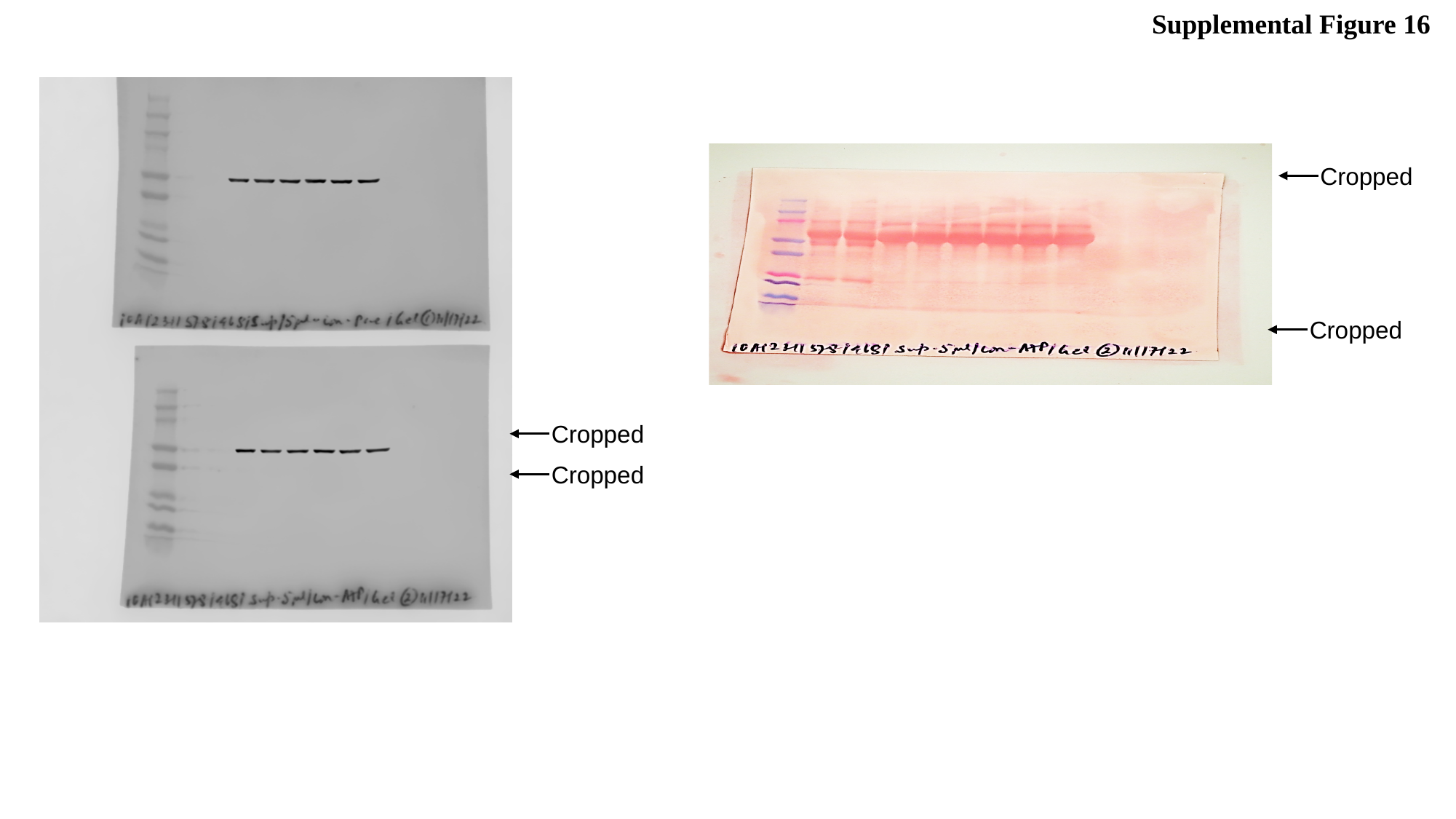

Supplemental Figure 16
 Cropped
 Cropped
 Cropped
 Cropped

### Slide 17
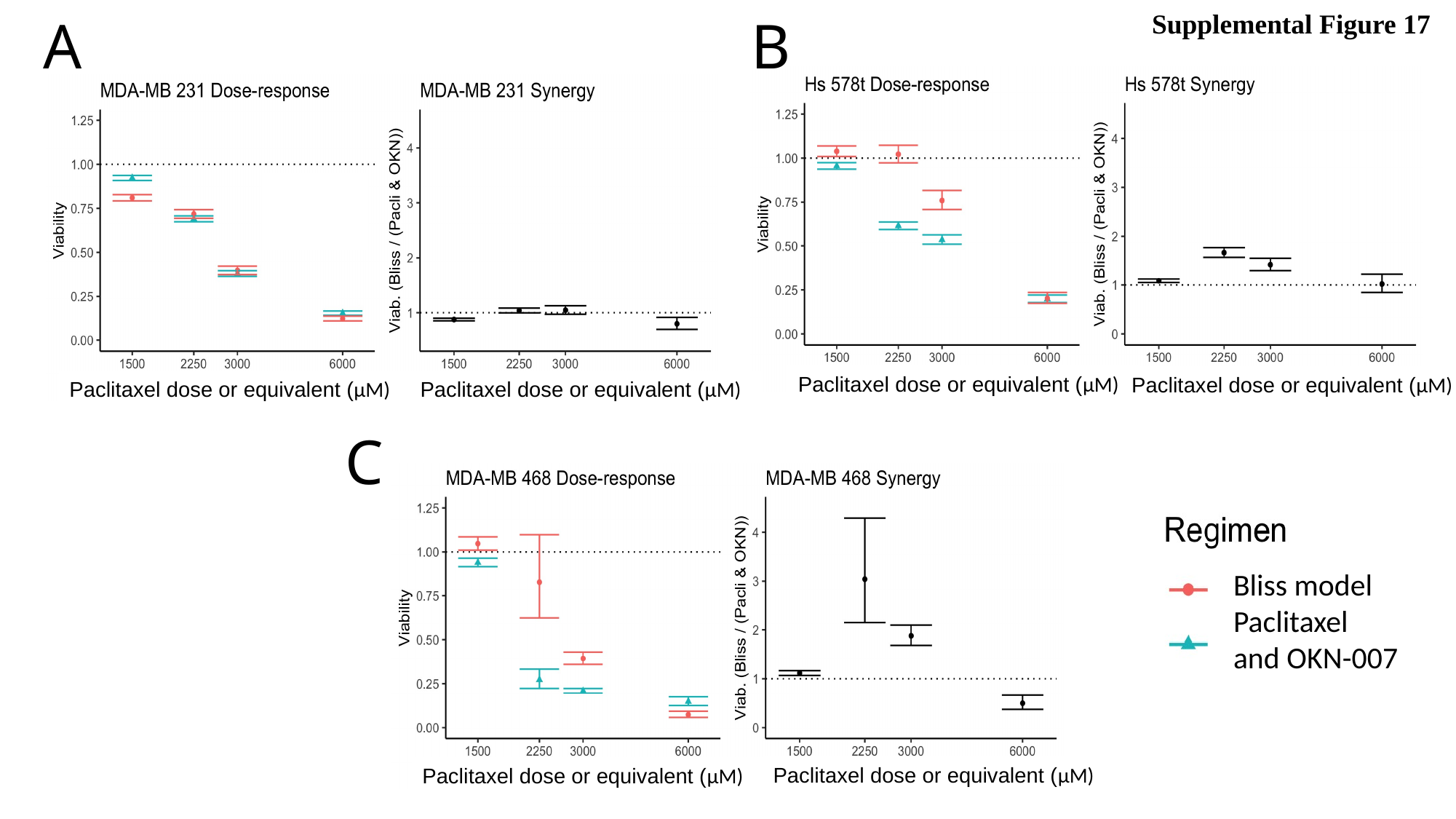

Supplemental Figure 17
A
B
Paclitaxel dose or equivalent (µM)
Paclitaxel dose or equivalent (µM)
Paclitaxel dose or equivalent (µM)
Paclitaxel dose or equivalent (µM)
C
Bliss model
Paclitaxel and OKN-007
Paclitaxel dose or equivalent (µM)
Paclitaxel dose or equivalent (µM)

### Slide 18
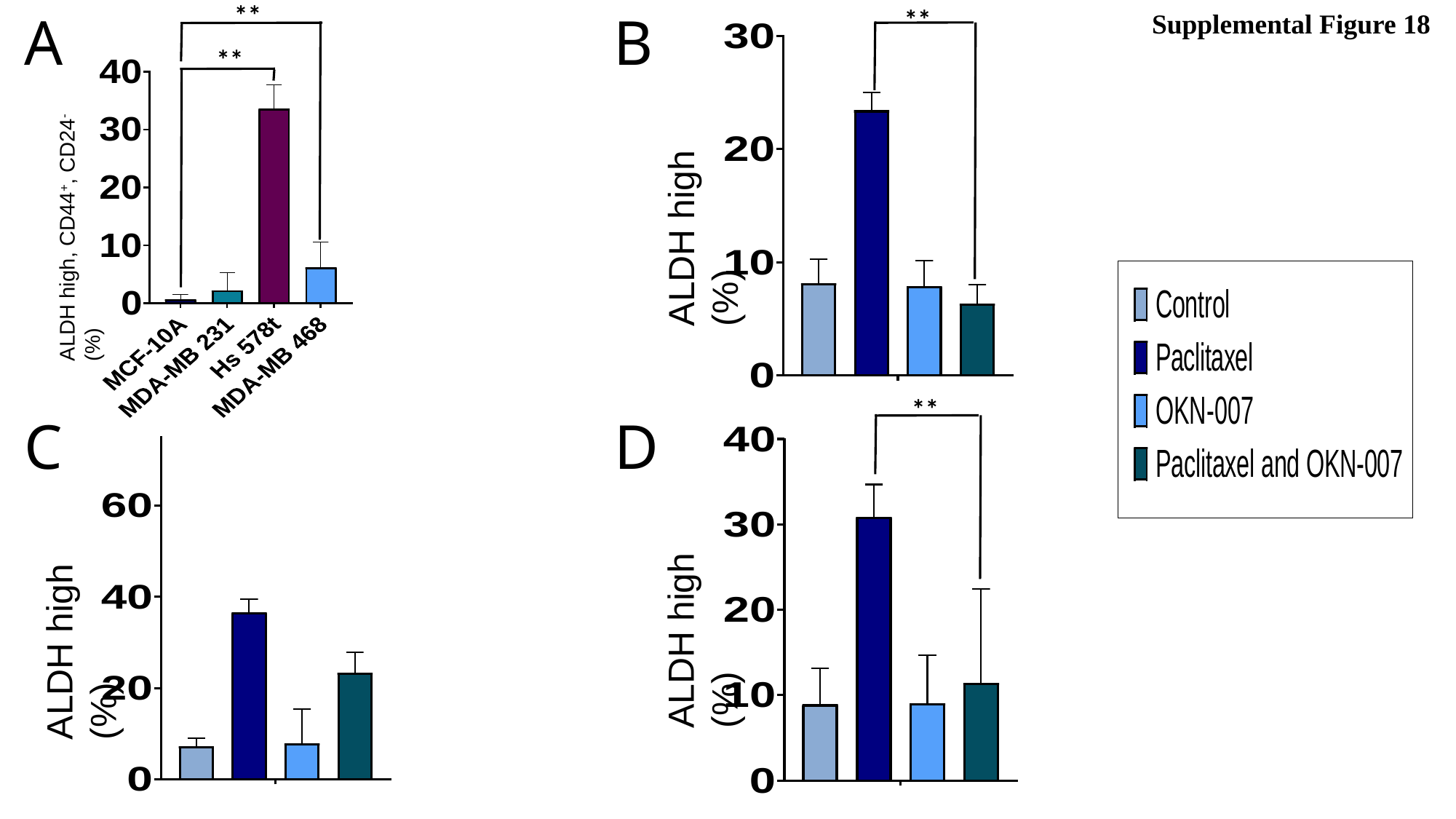

**
**
B
A
Supplemental Figure 18
**
ALDH high, CD44^+^, CD24^-^ (%)
ALDH high (%)
**
C
D
ALDH high (%)
ALDH high (%)

### Slide 19
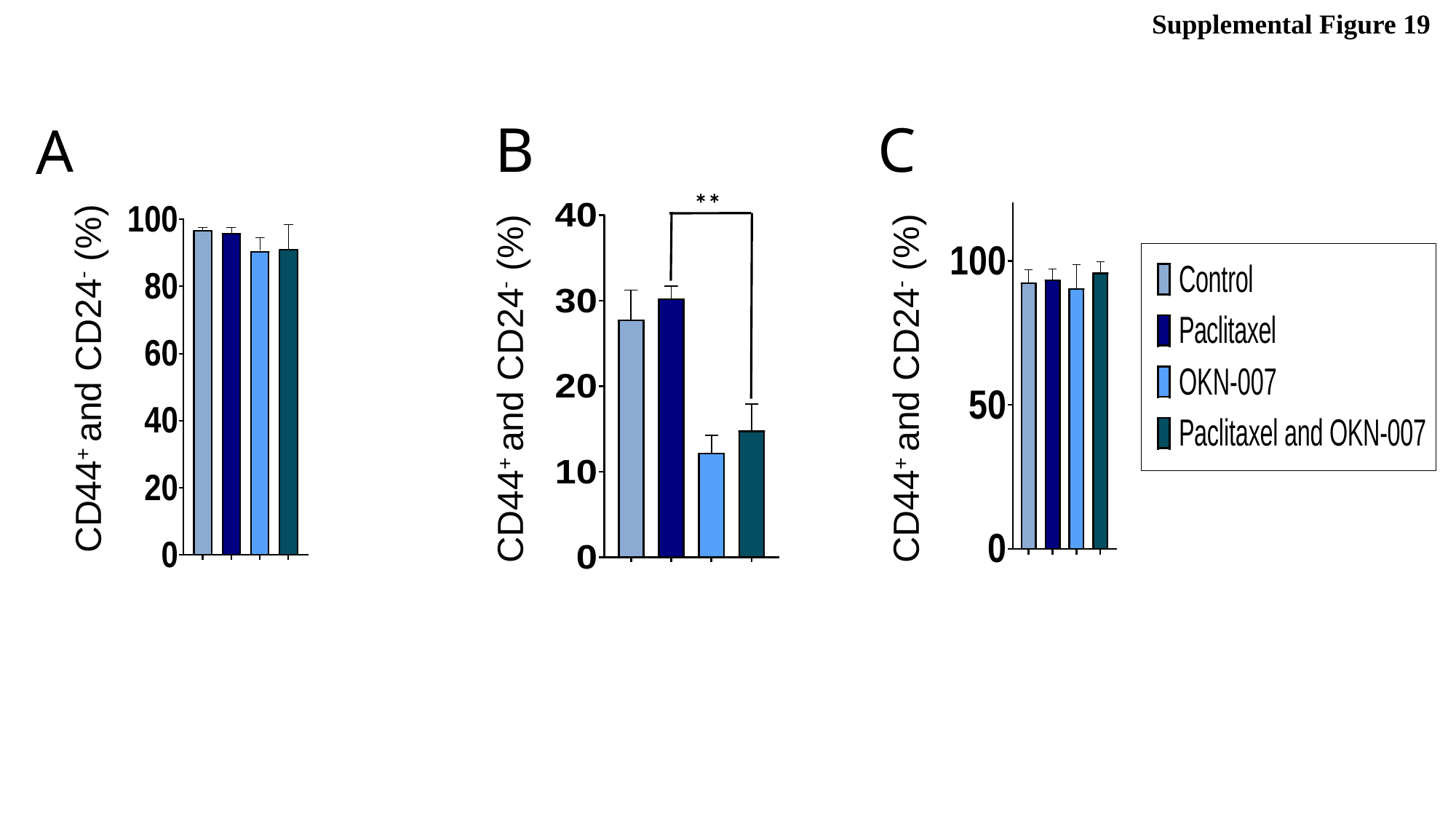

Supplemental Figure 19
B
C
A
CD44^+^ and CD24^-^ (%)
**
CD44^+^ and CD24^-^ (%)
CD44^+^ and CD24^-^ (%)

### Slide 20
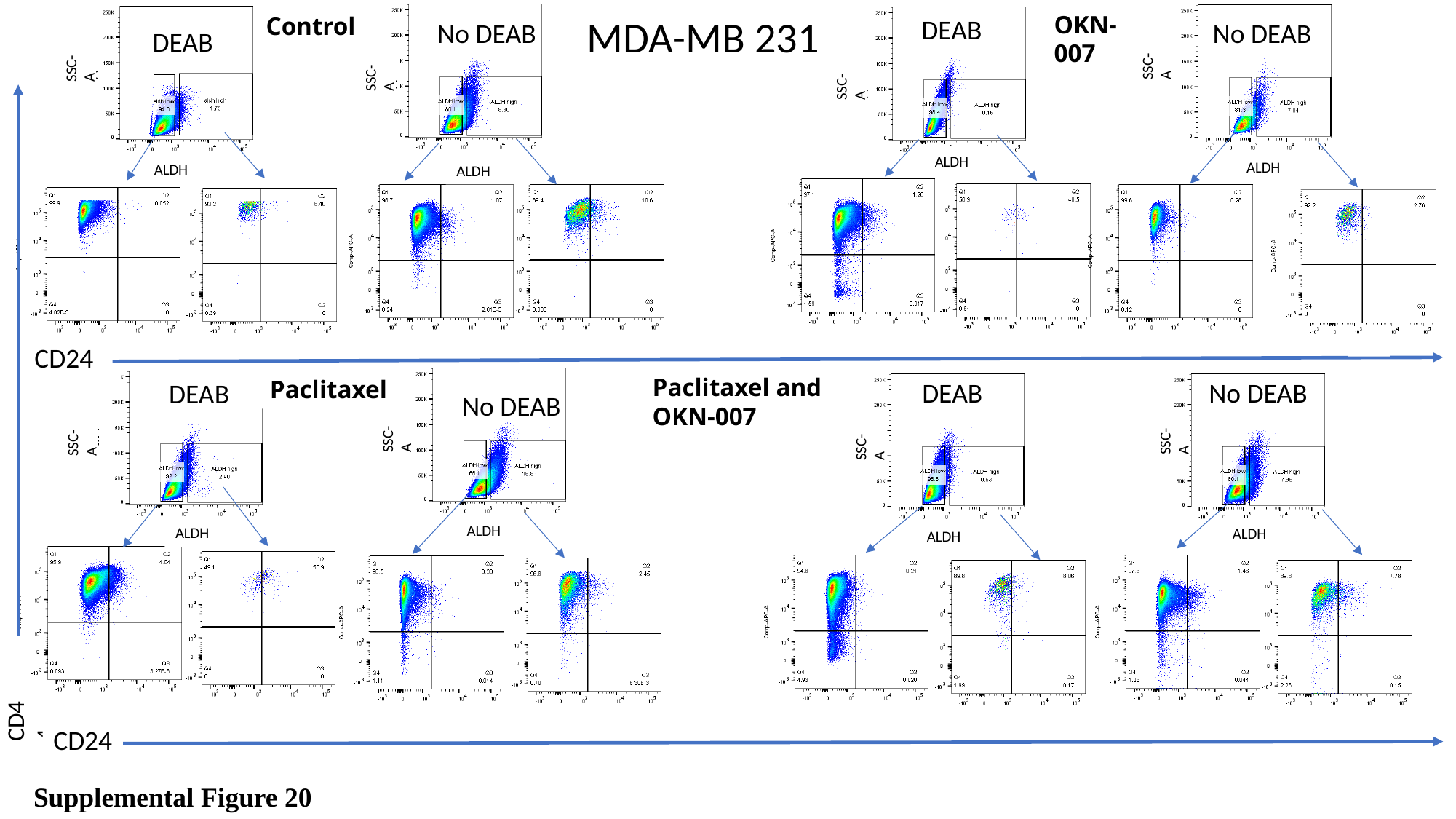

OKN-007
MDA-MB 231
Control
DEAB
No DEAB
No DEAB
DEAB
SSC-A
SSC-A
SSC-A
SSC-A
SSC-A
SSC-A
SSC-A
SSC-A
SSC-A
ALDH
ALDH
ALDH
ALDH
ALDH
ALDH
ALDH
ALDH
CD24
Paclitaxel and OKN-007
Paclitaxel
DEAB
No DEAB
DEAB
No DEAB
SSC-A
SSC-A
SSC-A
SSC-A
ALDH
ALDH
ALDH
ALDH
CD44
CD24
Supplemental Figure 20

### Slide 21
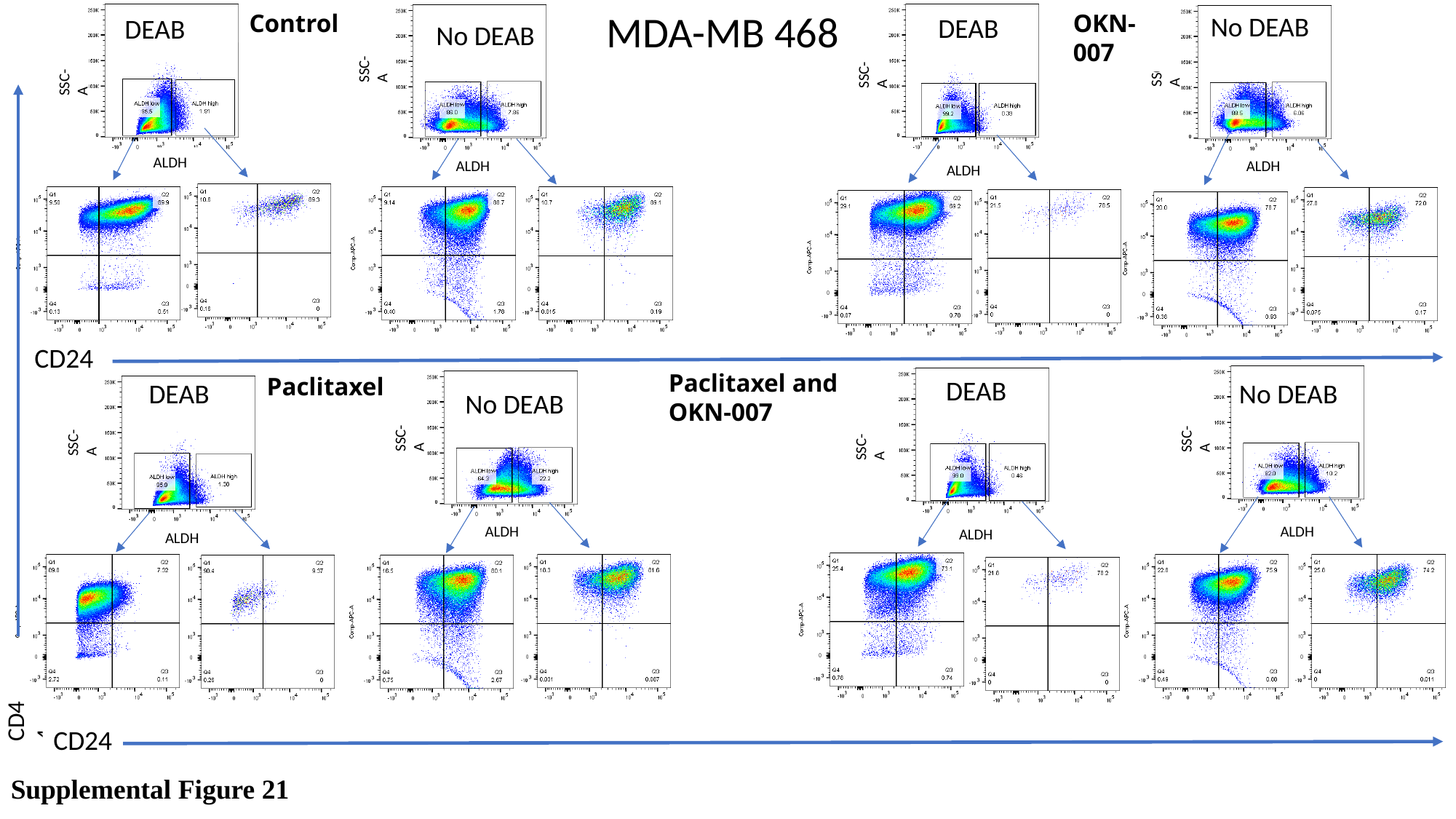

MDA-MB 468
Control
OKN-007
No DEAB
DEAB
DEAB
No DEAB
SSC-A
SSC-A
SSC-A
SSC-A
ALDH
ALDH
ALDH
ALDH
CD24
Paclitaxel and OKN-007
Paclitaxel
DEAB
DEAB
No DEAB
No DEAB
SSC-A
SSC-A
SSC-A
SSC-A
ALDH
ALDH
ALDH
ALDH
CD44
CD24
Supplemental Figure 21

### Slide 22
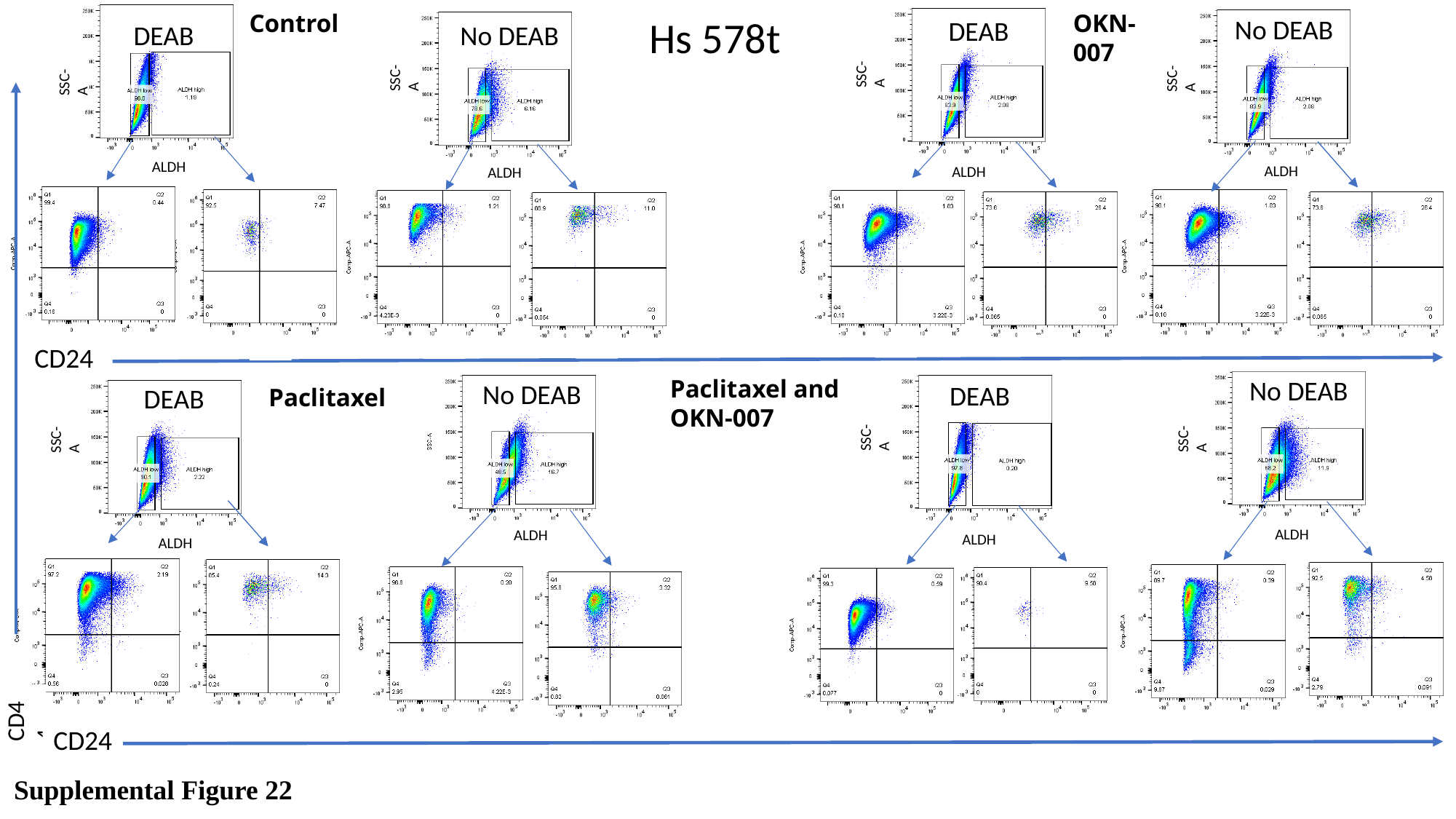

Control
OKN-007
Hs 578t
No DEAB
DEAB
No DEAB
DEAB
SSC-A
SSC-A
SSC-A
SSC-A
ALDH
ALDH
ALDH
ALDH
ALDH
CD24
No DEAB
Paclitaxel and OKN-007
No DEAB
DEAB
DEAB
Paclitaxel
SSC-A
SSC-A
SSC-A
ALDH
ALDH
ALDH
ALDH
CD44
CD24
Supplemental Figure 22

### Slide 23
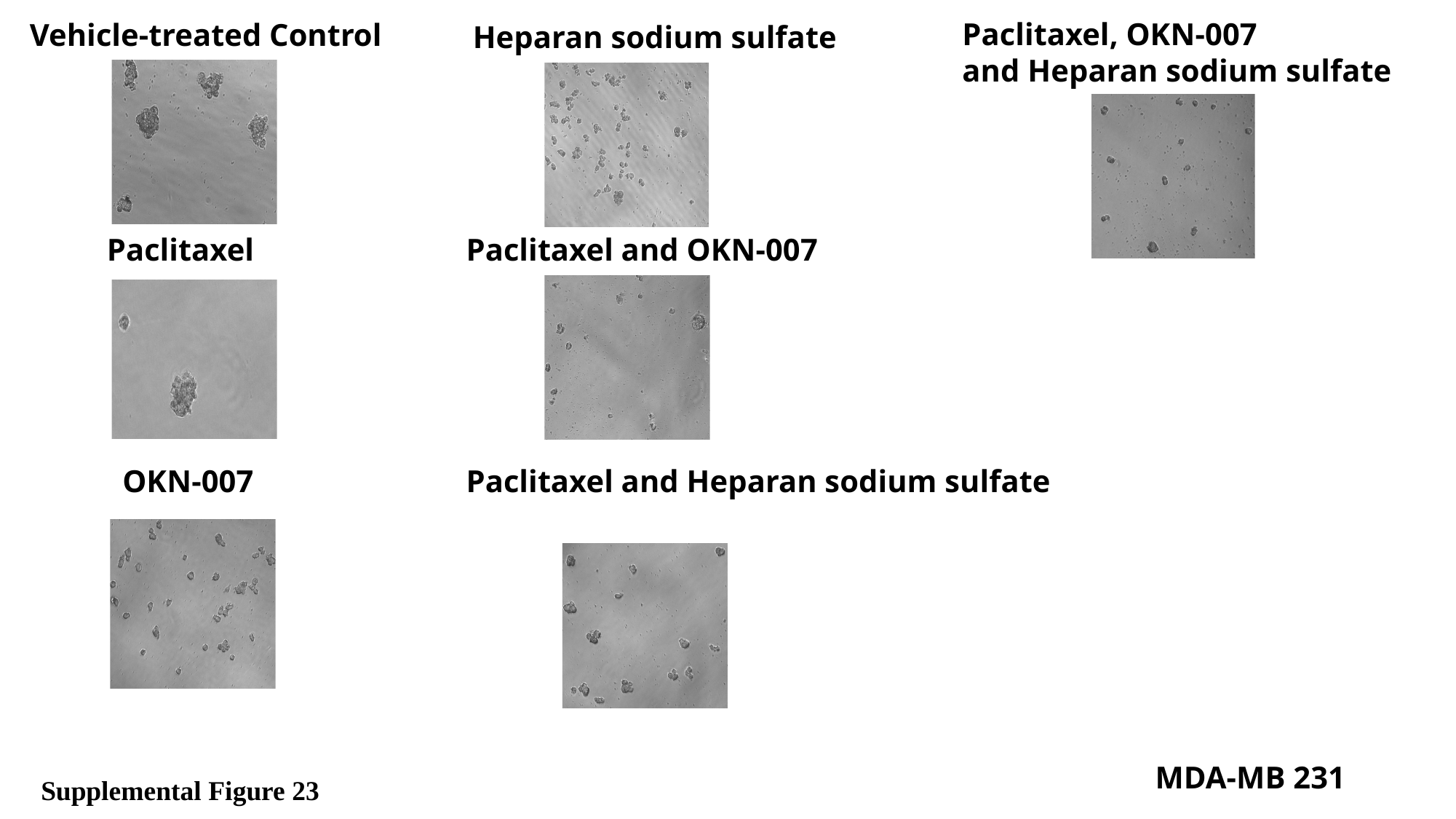

Paclitaxel, OKN-007
and Heparan sodium sulfate
Vehicle-treated Control
Heparan sodium sulfate
Paclitaxel
Paclitaxel and OKN-007
OKN-007
Paclitaxel and Heparan sodium sulfate
MDA-MB 231
Supplemental Figure 23

### Slide 24
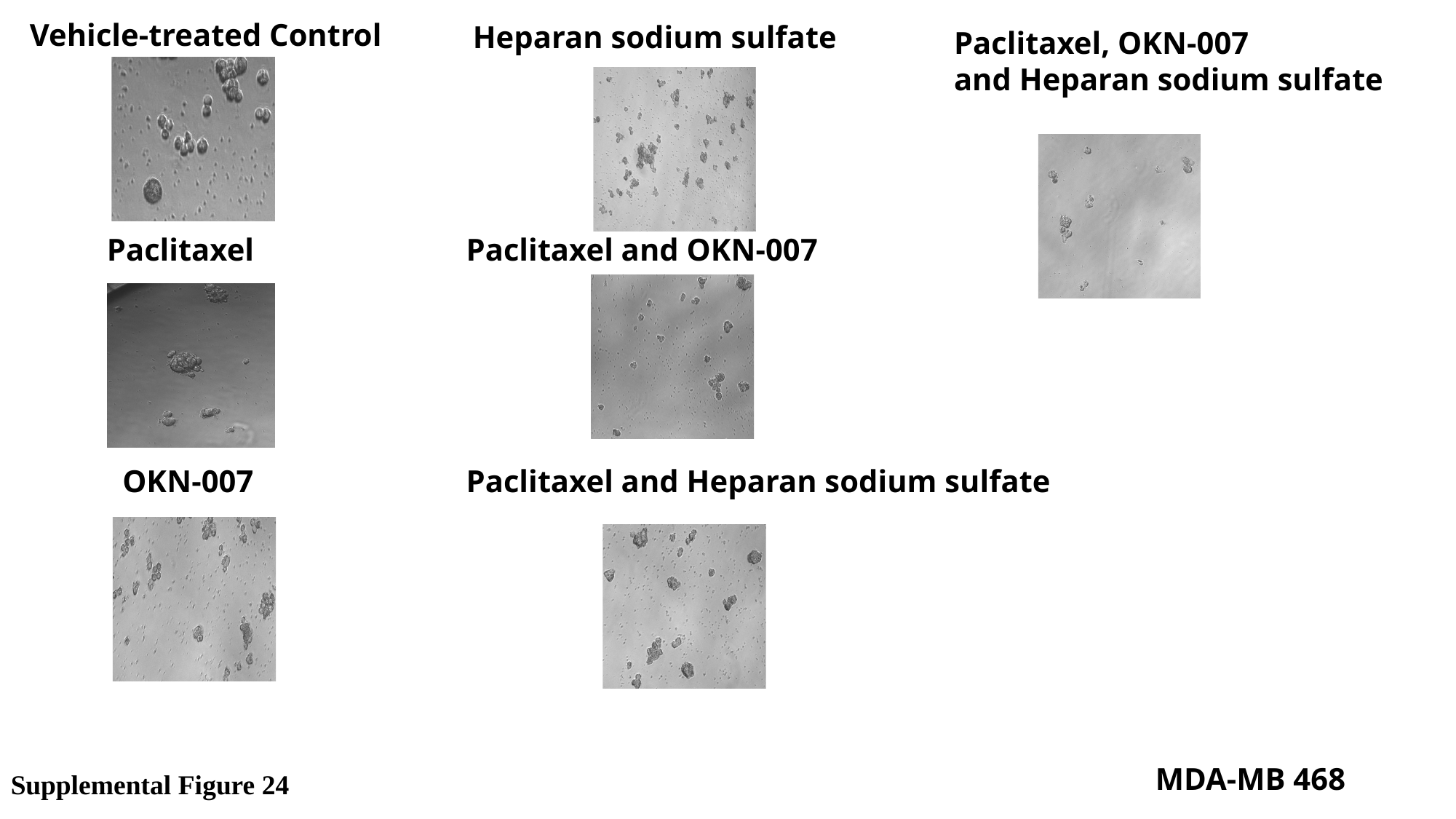

Vehicle-treated Control
Heparan sodium sulfate
Paclitaxel, OKN-007
and Heparan sodium sulfate
Paclitaxel
Paclitaxel and OKN-007
OKN-007
Paclitaxel and Heparan sodium sulfate
MDA-MB 468
Supplemental Figure 24

### Slide 25
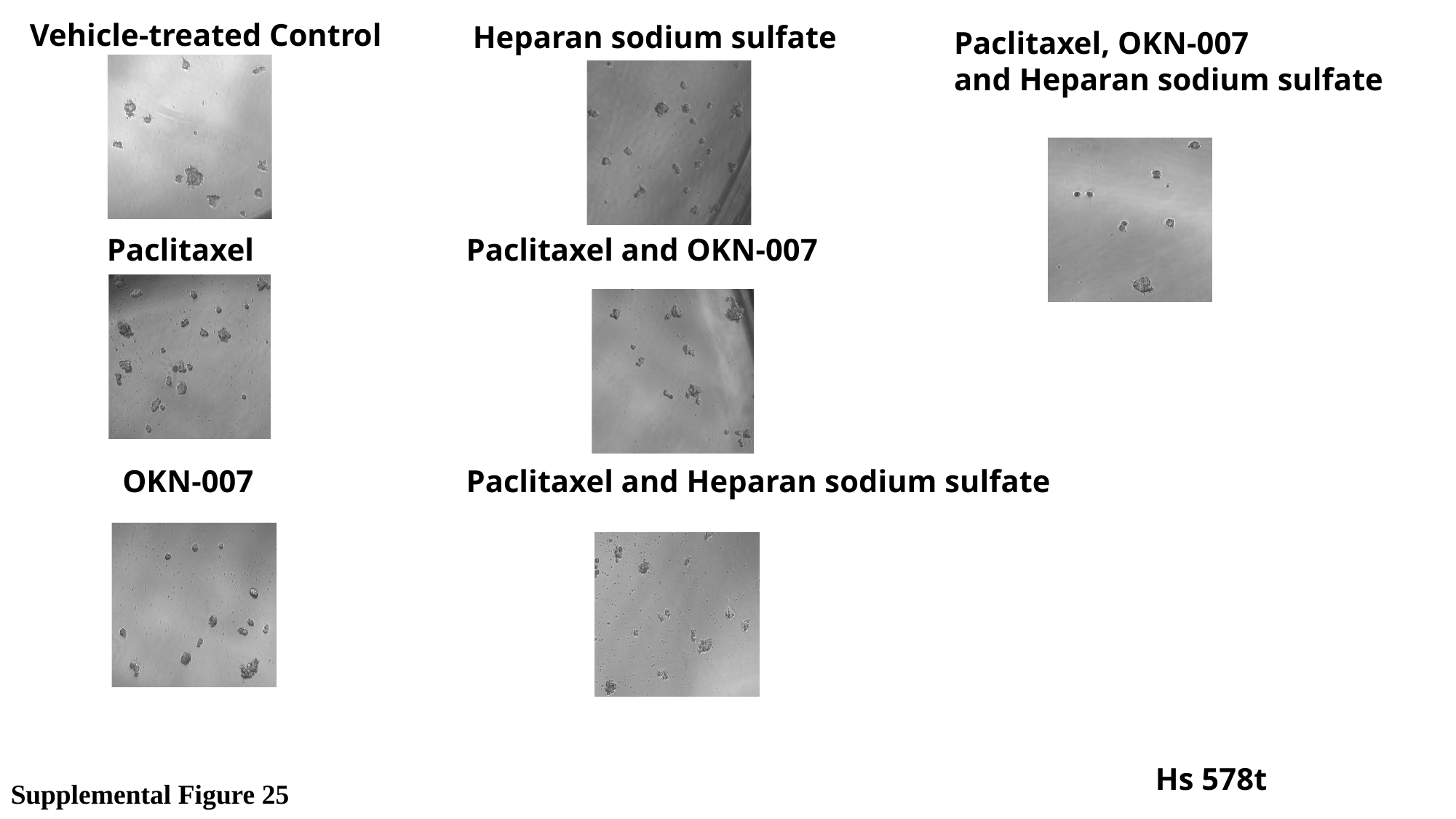

Vehicle-treated Control
Heparan sodium sulfate
Paclitaxel, OKN-007
and Heparan sodium sulfate
Paclitaxel
Paclitaxel and OKN-007
OKN-007
Paclitaxel and Heparan sodium sulfate
Hs 578t
Supplemental Figure 25
