## Supplemental Figure Legends for "Sulfatase 2 Inhibition Sensitizes Triple-Negative Breast Cancer Cells to Chemotherapy Through Augmentation of Extracellular ATP"

**Supplemental Figure 1: Sulfatase expression analysis.** Expression analysis of **(A)** sulfatase 1 **(B)** sulfatase 2 protein at different stages of breast cancer. **(C)** Messenger RNA expression analysis of sulfatases in TNBC MDA-MB 231, Hs 578t and MDA-MB 468 cell lines. This data was obtained from Expression Atlas and TSVdb.

**Supplemental Figure 2:** **Full western blot for sulfatase 2 from Figure 3A. (A)**The full, un-cropped western blot for sulfatase 2 from Figure 3A. **(B)** The full, un-cropped Ponceau S stain image for sulfatase 2 from Figure 3A.

**Supplemental Figure 3:** **Full western blot for sulfatase 2 from Figure 3B.** The full, un-cropped western blot for sulfatase 2 from Figure 3B.

**Supplemental Figure 4:** **Full western blot for sulfatase 2 from Figure 3C.** The full, uncropped western blot for **(A)** sulfatase 2 and **(B)** GAPDH from Figure 3C.

**Supplemental Figure 5:** **Full western blot for sulfatase 1 from Figure 4A. (A)**The full, uncropped western blot for sulfatase 1 from Figure 3A. **(B)** The full, uncropped Ponceau S stain image for sulfatase 1 from Figure 4A.

**Supplemental Figure 6:** **Full western blot for sulfatase 1 from Figure 4B.** The full, uncropped western blot for sulfatase 1 from Figure 4B.

**Supplemental Figure 7:** **Full western blot for sulfatase 1 from Figure 4C.** The full, uncropped western blot for **(A)** sulfatase 1 and **(B)** GAPDH from Figure 4C.

**Supplemental Figure 8: ELISA analysis of sulfatases.** ELISAs were performed to examine the basal levels of **(A)** sulfatase 2 and **(B)** sulfatase 1 in supernatants of TNBC MDA-MB 231, Hs 578t and MDA-MB 468 cell lines and non-tumorigenic immortal epithelial mammary MCF-10A cells. MCF-10A expressed more sulfatase 1 in comparison to the TNBCs while TNBCs expressed more sulfatase 2 in comparison to MCF-10A cells. Standard deviation was calculated from three independent experiments performed in triplicate. The student’s t-test was performed to determine significance with * representing p<0.05 and ** representing p<0.01 comparing the protein expression MCF-10A to the protein expressions of TNBC cell lines.

**Supplemental Figure 9:** **Flow cytometry analysis of sulfatase 1 and 2 in TNBC and MCF-10A cells.** **(A)** Cell surface basal expression of sulfatase 1 was examined in TNBC and MCF-10A cells via flow cytometry; sulfatase 1 was expressed at higher levels in the MCF-10A cells than TNBC cells. The standard deviation was calculated from three independent experiments performed in triplicate. The student’s t-test was performed to determine significance with * representing p<0.05 and ** representing p<0.01 comparing the protein expression in MCF-10A to the protein expressions in TNBC cell lines. **(B)** Histogram obtained from flow cytometry analysis of the cell surface expression of sulfatase 2 of the TNBC and MCF-10A cell lines.

**Supplemental Figure 10: Immunohistochemistry for sulfatase 2 staining. (A)** Normal breast tissue on slides (2) stained for sulfatase 2. **(B)** Ductal carcinoma in situ (DCIS) tissue on slides (3) stained for sulfatase 2. **(C)** Slide key for the breast cancer tissue array orientation.

**Supplemental Figure 11: Statistical analysis for sulfatase 2 amongst normal tissue, DCIS, and various grades of cancer. (A)** Pairwise comparisons using Dunn’s test showed that there was no significant difference between the average percentages of cells showing any level of staining for sulfatase 2 in the tissue sections of normal tissue, DCIS, and various grades of cancer**. (B)** Kruskal-Wallis test demonstrated that there was no significant difference between the average percentages of cells showing any level of staining for sulfatase 2 in the tissue sections of normal tissue, DCIS, and various grades of cancer. **(C)** Pairwise comparisons using Dunn’s test indicated that there was a significant difference between percentages of cells staining moderately for sulfatase 2 in tissue sections of cancer grades 1 and 3 (p = 0.0127) with Grade 3 expressing more sulfatase 2. No other differences were statistically significant. **(D)** Pairwise comparisons using Dunn’s test indicated that there was a significant difference between the percentages of cells staining strongly for sulfatase 2 in tissue sections of cancer grades 1 and 3 (p = 0.0207) with grade 3 expressing more sulfatase 2 than grade 1. **(E)** Pairwise comparisons using Dunn’s test indicated that there was no significant difference between the percentages of cells in the tissue sections of normal tissue, DCIS, and various grades of cancer that were negative for staining for sulfatase 2 between.

**Supplemental Figure 12:** **Additional statistical analysis for sulfatase 2 staining amongst different progesterone receptor expression levels. (A)** Kruskal-Wallis test demonstrated that there was no significant difference between the average percentages of cells with any level of staining for sulfatase 2 in tissue sections of breast cancers with different progesterone receptor (PR) expression levels. **(B)** The Kruskal-Wallis test demonstrated that there was no significant difference between the average percentages of cells in tissue sections staining weakly for sulfatase 2 amongst breast cancers expressing different PR levels. **(C)** Pairwise comparisons using Dunn’s test showed that there was no significant difference between the average percentages of cells with moderate staining for sulfatase 2 amongst breast cancers with different PR expression levels. **(D)** Pairwise comparisons using Dunn’s test demonstrated that there was no significant difference between the percentages of cells staining strongly for sulfatase 2 in tissue sections of breast cancers that expressed different PR levels. **(E)** Kruskal-Wallis test demonstrated that there was no significant difference between the percentages of cells that stained negatively for sulfatase 2 in tissue section of breast cancers with different PR expression levels.

**Supplemental Figure 13:** **Additional statistical analysis for sulfatase 2 staining and comparing % Ki67 expression levels. (A)** Pairwise comparisons using Dunn’s test indicated that there was a significant difference between the percentages of cells expressing any level of sulfatase 2 in tissue sections of cancers with different levels of Ki67 expression. The higher % Ki67 expressed more sulfatase 2. **(B)** Pairwise comparisons using Dunn’s test indicated that there was a significant difference between the percentages of cells with weak positive staining for sulfatase 2 in tissue sections of cancers with differing Ki67 expression levels. The higher % Ki67 expressed more sulfatase 2. **(C)** Pairwise comparisons using Dunn’s test indicated that there was a significant difference between the percentage of cells with moderately positive staining for sulfatase 2 in tissue sections of cancers with differing Ki67 expression levels. The higher % Ki67 expressed more sulfatase 2.: **(D)** Pairwise comparisons using Dunn’s test indicated that there was a significant difference between the percentages of cells staining strongly positive for sulfatase 2 in tissue sections of cancers with differing Ki67 expression levels. The higher % Ki67 expressed more sulfatase 2 **(E)** Pairwise comparisons using Dunn’s test indicated that there was a significant difference between the percentages of cells that were negative for sulfatase 2 staining in tissue sections of cancers with differing Ki67 expression levels.

**Supplemental Figure 14:** **Western blot analysis for TNBC and MCF-10A cells treated with paclitaxel or ATP**. TNBC MDA-MB 231, Hs 578t and MDA-MB 468 cells and nontumorigenic immortal mammary epithelial MCF-10A cells were treated with **(A)** paclitaxel (100 µM) for 6 hours or **(B)** ATP (500 µM) for 48 hours and 5 μl cell supernatants were probed sulfatase 2. Ponceau S staining was performed immediately after transfer onto the nitrocellulose membranes to demonstrate loading of the proteins. Similar results were obtained in biological replicate experiments.

**Supplemental Figure 15:** **Full western blot for sulfatase 2 from Supplemental Figure 14A. (A)** The full, un-cropped western blot for sulfatase 2 from Supplemental Figure 14A. **(B)** The full, un-cropped Ponceau S stain image for sulfatase 2 from Supplemental Figure 14A.

**Supplemental Figure 16:** **Full western blot for sulfatase 2 from Supplemental Figure 14B. (A)** The full, un-cropped western blot for sulfatase 2 from Supplemental Figure 14B. **(B)** The full, un-cropped Ponceau S stain image for sulfatase 2 from Supplemental Figure 14B.

**Supplemental Figure 17:** **Statistical analysis of treated cells with sulfatase inhibitor OKN-007 and chemotherapeutic agent paclitaxel.** Dose response and synergy graphs are displayed for TNBC **(A)** MDA-MB 231 **(B)** Hs 578t and **(C)** MDA-MB 468 with increasing concentrations of paclitaxel, OKN-007 or the co-treatment administered for 48 hours. For the dose response graphs, the bliss model (orange) and the co-treatment (paclitaxel and OKN-007) are shown. There is some synergy (<0.1-1.0) for some dose combinations for MDA-MB 231 cells while there were some drug dose combinations that were additive (1-1.2) for Hs 578t and MDA-MB 468 cells. These graphs were obtained from three independent experiments performed in triplicate.

**Supplemental Figure 18:** **ALDH expression in TNBC cells.** The percent of cells is shown for **(A)** MDA-MB 231 **(B)** MDA-MB 468 and **(C)** Hs 578t that are considered ALDH high. TNBCs treated with paclitaxel alone produced the most cells that expressed ALDH at high concentrations. Standard deviation was calculated from three independent experiments performed in triplicate. 1-way ANOVA with Tukey’s HSD was applied to ascertain significance. * represents p<0.05 and ** represents p<0.01 when comparing paclitaxel to paclitaxel and OKN-007.

**Supplemental Figure 19: CD44 and CD24 expressions in TNBC cells.** The percent of cells is shown for **(A)** MDA-MB 231 **(B)** MDA-MB 468 and **(C)** Hs 578t that are considered CD44 positive and CD24 negative. MDA-MB 231 and Hs 578t cells highly express CD44 under all drug conditions; whereas, MDA-MB 468 cells treated with vehicle or paclitaxel expressed more CD44 than those treated with OKN-007 or the co-treatment of OKN-007 and paclitaxel. Standard deviation was calculated from three independent experiments performed in triplicate. 1-way ANOVA with Tukey’s HSD was applied to ascertain significance. * represents p<0.05 and ** represents p<0.01 when comparing paclitaxel to paclitaxel and OKN-007.

**Supplemental Figure 20:** **Cancer-initiating cell dot plots for OKN-007 and paclitaxel-treated MDA-MB 231 cells**. Dot plots examining the cancer-initiating cells that are ALDH high, CD44 positive, and CD24 negative are presented for treated MDA-MB 231 cells with paclitaxel, sulfatase inhibitor OKN-007, or both drug agents. Diethylaminobenzaldehyde (DEAB) is a specific inhibitor of ALDH, that can be used as a background fluorescence control. Three independent experiments were performed in triplicate.

**Supplemental Figure 21:** **Cancer-initiating cell dot plots for OKN-007 and paclitaxel-treated Hs 578t cells**. Dot plots examining the cancer-initiating cells that are ALDH high, CD44 positive, and CD24 negative are presented for treated Hs 578t cells with paclitaxel, sulfatase inhibitor OKN-007, or both drug agents. DEAB is used as a background fluorescence control. Three independent experiments were performed in triplicate.

**Supplemental Figure 22:** **Cancer-initiating cell dot plots for OKN-007 and paclitaxel-treated MDA-MB 468**. Dot plots examining the cancer-initiating cells that are ALDH high, CD44 positive, and CD24 negative are presented for treated MDA-MB 468 cells with paclitaxel, sulfatase inhibitor OKN-007, or both drug agents. DEAB is used as a background fluorescence control. Three independent experiments were performed in triplicate.

**Supplemental Figure 23:** **Tumorsphere efficiency assay images for treated MDA-MB 231 cells**. Tumorsphere images obtained from the Etaluma™ Lumascope 620 (10X) are displayed for each treatment of MDA-MB 231 cells with paclitaxel, OKN-007, heparan sodium sulfate, or the different combinations. Three independent experiments were performed in triplicate.

**Supplemental Figure 24:** **Tumorsphere efficiency assay images for treated MDA-MB 468 cells**. Tumorsphere images obtained from the Etaluma™ Lumascope 620 (10X) are displayed for each treatment of MDA-MB 468 cells with paclitaxel, OKN-007, heparan sodium sulfate, or the different combinations. Three independent experiments were performed in triplicate.

**Supplemental Figure 25:** **Tumorsphere efficiency assay images for treated Hs 578t cells**. Tumorsphere images obtained from the Etaluma™ Lumascope 620 (10X) are displayed for each treatment of Hs 578t cells with paclitaxel, OKN-007, heparan sodium sulfate, or the different combinations. Three independent experiments were performed in triplicate.
